# 1RS.1BL molecular resolution provides novel contributions to wheat improvement

**DOI:** 10.1101/2020.09.14.295733

**Authors:** Zhengang Ru, Angela Juhasz, Danping Li, Pingchuan Deng, Jing Zhao, Lifeng Gao, Kai Wang, Gabriel Keeble-Gagnere, Zujun Yang, Guangrong Li, Daowen Wang, Utpal Bose, Michelle Colgrave, Chuizheng Kong, Guangyao Zhao, Xueyong Zhang, Xu Liu, Guoqing Cui, Yuquan Wang, Zhipeng Niu, Liang Wu, Dangqun Cui, Jizeng Jia, Rudi Appels, Xiuying Kong

## Abstract

Wheat-rye 1RS.1BL translocation has a significant impact on wheat yield and hence food production globally. However, the genomic basis of its contributions to wheat improvement is undetermined. Here, we generated a high-quality assembly of 1RS.1BL translocation comprising 748,715,293 bp with 4,996 predicted protein-coding genes. We found the size of 1RS is larger than 1BS with the active centromere domains shifted to the 1RS side instead of the 1BL side in Aikang58 (AK58). The gene alignment showed excellent synteny with 1BS from wheat and genes from 1RS were expressed well in wheat especially for 1RS where expression was higher than that of 1BS for the grain-20DPA stage associated with greater grain weight and negative flour quality attributes. A formin-like-domain protein FH14 (*TraesAK58CH1B01G010700*) was important in regulating cell division. Two PPR genes were most likely the genes for the multi fertility restoration locus *Rf ^multi^*. Our data not only provide the high-resolution structure and gene complement for the 1RS.1BL translocation, but also defined targets for enhancing grain yield, biotic and abiotic stress, and fertility restoration in wheat.

## INTRODUCTION

The 1RS.1BL translocation chromosome was one of the earliest of so-called alien chromatin additions into wheat and is generally considered to be associated with the disease resistance (Zeller, 1973) and a step-change in yield achieved with the release of the Veery lines by CIMMYT (Rajaram et al., 1983). A survey by R Schlegel showed that approximately 30% of wheat cultivars released after the year 2000 carry the 1RS.1BL translocation (http://www.rye-gene-map.de/rye-introgression/) (Schlegel and Korzun, 1997). The *Lr26, Sr31, Yr9, Sr50* rust resistance genes and the powdery mildew, *Mlg* locus have been identified on 1RS.1BL chromosomes as well as genetic factors affecting root biomass (Mago et al., 2002 and 2015; Ehdaie et al., 2003; Waines and Ehdaie, 2007; Sharma et al., 2011). The grain yield associated with 1RS.1BL has been shown to be disrupted by 1RS-1BS recombinants in the terminal region of 1RS as characterized in field trials of wheat accessions carrying these recombinants in the same genetic background (Lukaszewski et al., 2000; Howell et al., 2014 and 2019). The 1RS.1BL chromosome has also been factored into maintaining male sterility in hybrid wheat programs (Lukaszewski et al., 2017).

The 1RS rye chromosome segment in wheat varieties has at least 3 sources, namely, as 1RS.1BL in *Triticum aestivum* cv Salmon (Japan), as 1RS.1BL in *T. aestivum* varieties originating from Germany and as 1RS.1AL in *T. aestivum* cv Amigo from the USA (Schlegel and Korzun, 1997). The 1RS.1BL translocation has been widely introduced into wheat cultivars globally for conferring a broad-range of resistance to races of powdery mildew and rusts, environmental adaptability and yield performance (Zeller, 1973; Rajaram et al., 1983; Schlegel and Korzun, 1997; Bartoš and Bareš, 1971; Sukumaran et al., 2015).

At a structure level, 1RS.1BL was one of the early chromosomes to have specific DNA sequences assigned to it through the analysis of chromosomes using inbred lines of rye to facilitate locating molecular markers and agronomic traits to 1RS as well as the entire rye genome (Lawrence and Appels, 1986; Miedaner et al., 2012; Bauer et al., 2016). *In situ* hybridization technology with labelled sequenced probes has allowed the convenient microscopic visualization and identification of translocations involving 1RS (Appels et al., 1978; McIntyre et al., 1990; Liu et al., 2017).

The favorable agronomic features have been a significant driver for sequencing the 1RS.1BL chromosome in the China wheat cultivar, AK58. In the available rye genome data from an inbred rye line 27,784 gene models (and segments) were sourced for assigning gene models to the 1RS segment and these generally fell within gene models aligned from the wheat reference genome sequence (Bauer et al., 2016; IWGSC et al., 2018). This study provides the linear genome level structure of AK58-1RS.1BL utilizing a combination of Illumina and PacBio sequencing with de novo NR Magic for the initial assembly followed by HiC scaffolding and alignment to high density molecular genetic maps to generate the final assembly. The genome structure identified new gene models, several multi-gene families likely to be involved in yield attributes associated with 1RS.1BL and resolved gene families involved in disease resistance and other agronomic traits.

## RESULTS

### Assembly of the 1RS.1BL genome of wheat cv AK58

The details of the assembly are provided in Figure S1 and Methods, and include the de novo NR Magic software to carry out a primary alignment into contigs, followed by the HiC process for scaffolding the contigs. Finally, we generated a high-quality chromosome-scale assembly of 1RS.1BL translocation with a total length of 748,715,293 bp and predicted 4,996 genes using four different annotation pipelines and alignment with the IWGSC RefSeq v1.0 annotation. The alignment of our AK58_chr1RS.1BL_v6 to the reference Chinese Spring (CS) 1B is shown in Figure 1a. The AK58_chr1RS.1BL_v6 assembly was examined in detail in the terminal 22 Mb region because, in general, this region of wheat genome assemblies can be problematical with respect to contig orientation. The assembly shown was the best alignment to available genetic mapping information for 1RS.1BL, as shown in Figure 1b (Mago et al., 2002; Howell et al., 2014; Sharma et al., 2009), using the markers, gamma secalin (8.82 Mb) and BE444266 (24.41 Mb). The IB267 (0.815 Mb) and iag95 (3.92 Mb) markers were located following discussions with J Dubcovsky (pers. comm., see Figure 1b).

**Figure 1.**
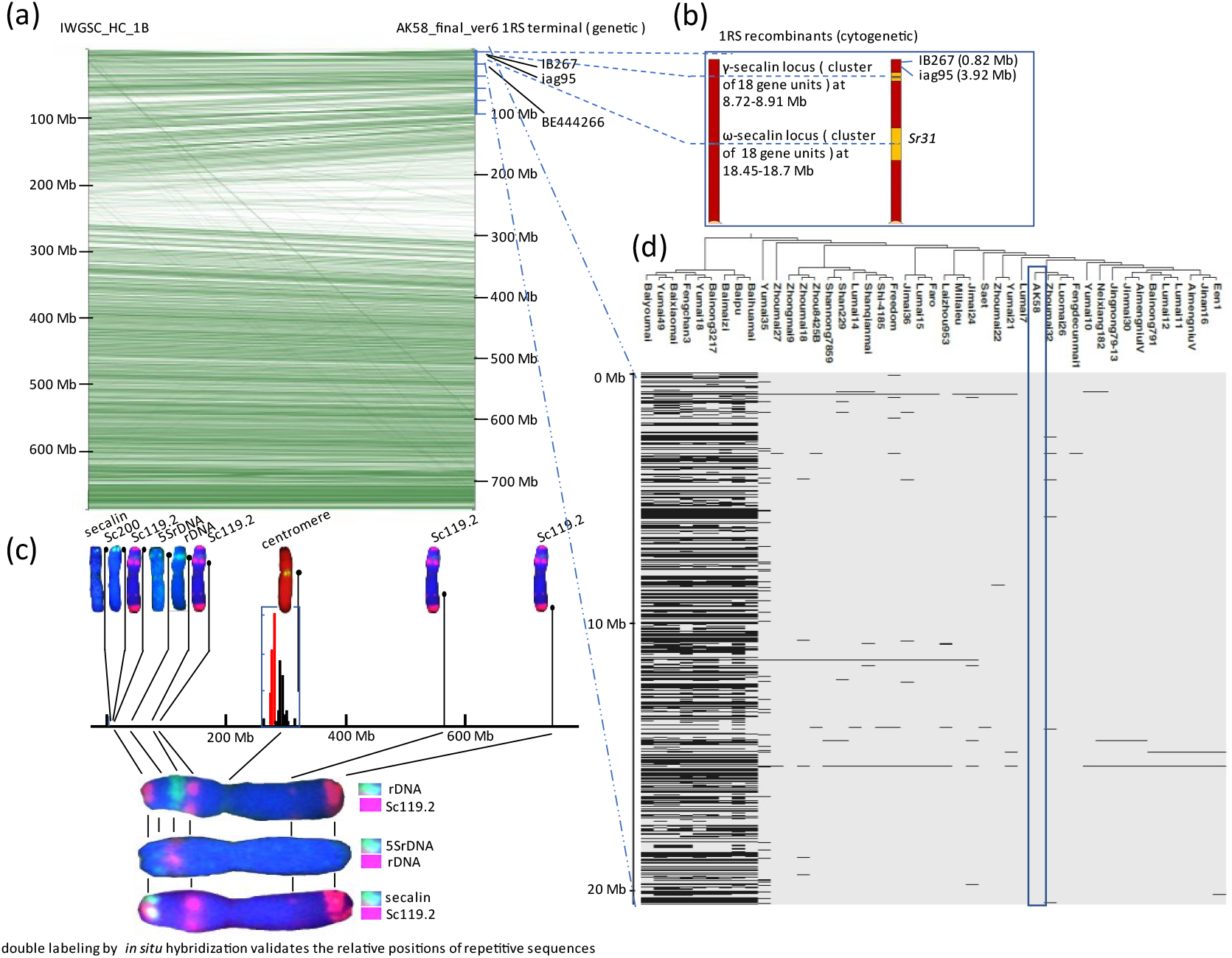
Structure of the AK58 1RS.1BL chromosome. (a) The high confidence (HC) gene models from the IWGSC-Refseq v1.0/-Refseq v2.0 assembly for chromosome 1B of CS wheat have been aligned with their syntenic partner gene models in AK58 1RS.1BL using open source Pretzel (IWGSC et al., 2018; Keeble-Gagnère et al., 2019). (b) Summary of available genetic and cytogenetic details for the terminal 22 Mb region of 1RS showing 1RS (red)-1BS (yellow) recombinants that disrupt the yield attributes associated with the 1RS.1BL chromosome (Howell et al., 2014 and 2019). (c) *In situ* hybridization of repetitive sequence probes typically used to identify rye chromosomes to allow a broad level validation of the 1RS assembly. (d) Distribution of SNPs along representative sections of the 1RS.1BL chromosomes based on a 660K SNP-chip analysis (Figure S2 and Table S1). Nine wheat cultivars (left most clusters) and 36 1RS containing cultivars (remaining columns in the Figure) where the dark horizontal lines indicate a SNP in the respective position that is different in AK58 reference genome (blue box). Absence of a dark line indicates the alternate allele is the same as that in AK58.

### Alignment and *in situ* cross-referencing of the AK58-1RS.1BL genome

The *in situ* probes are generally repetitive and although the number of repeats was clearly collapsed during the assembly process, all the regions aligned by *in situ* hybridization using double labelling could be assigned positions in the new assembly (McIntyre et al., 1990; Liu et al., 2017; Zhang et al., 2004). Importantly the macro-level structure of 1RS.1BL could be validated in this way (Figure 1c). The repetitive array of Sc119.2 sequences at 117.6 Mb (31 copies of the core 45 bp repeat unit) on 1RS were under-represented in the assembly, relative to the array at 0.4 Mb (662, 45 bp repeat units) based on comparing the *in situ* hybridization signals which indicated qualitatively similar signals (Figure 1c). The amplification of repetitive gene families in 1RS (Sc119.2, Sc200) in positions that were not in a syntenic order has occurred against a background of a conserved syntenic order of high confidence (HC) gene models (Figure 1a). The repetitive gamma-secalin and omega-secalin gene families were located in syntenic positions relative to 1BS of CS. In Figure 1d, a portion of a diversity analysis is shown using a 660K SNP-chip for SNPs that could be clearly scored in 36 1RS containing lines (identified using the gamma-secalin based PCR probes) (Figure S2 and Table S1). A subset of 9 wheat lines are shown for comparison to confirm that at the macro-level the 1RS is a large haplotype block, as described by Cheng et al. (Cheng et al., 2019). At a micro-level at least 6 groups, or haplotypes, of 1RS.1BL could be identified using the AK58-1RS as a reference, and these are accounted for by considering the different rye genome sources used in the intense breeding efforts in China combining 1RS.1BL containing wheat lines in crosses with triticales, rye and alternative sources of 1RS.1BL (Figure 1d, Figure S2 and Table S1). The 1RS in AK58 groups with only 3 other 1RS lines.

In our assembly, the size of 1RS is 275 Mb, 28% larger than 1BS (215 Mb) and is consistent with the overall genome size of rye (7-8 Gb) being approximately one-third larger than the diploid genome of barley or wheat progenitors. A comparison of the TE-complement between 1RS, 1AS, and 1DS in AK58 and 1BS in CS indicated that 12 TE subfamilies (Figure S3 and Table S2) were dominant in 1RS with a total length in excess of 41.9 Mb (15.27% of the 1RS length), compared to only 4.9 Mb in 1AS, 3.5 Mb in 1DS and 4.8 Mb in 1BS of CS. The 12 rye dominant TE included nine LTRs. There were five TE families, *LTR-Gypsy-RLG_famc9.2, LTR-RLX_famc7, LTR-RLX famc21, Unknown-XXX_famc9,* and *Unknown-XXX_famc81,* in which the length ratios of 1RS/1AS, 1RS/1BS and 1RS/1DS are range from 4 to 203 (Table S2). Although most of the rye dominant TEs were distributed in 1RS, including gene-flanking regions, some were more prominent in the centromere region (LTR-Gypsy-RLG_famc36, Figure S3; see also centromere section below).

### Centromere structure at the 1RS-1BL boundary

The availability of a rye centromere sequence (pAWRC, Francki, 2001) that could be distinguished from the wheat centromere repeats, CRWs, allowed a more detailed analysis of the rye-wheat hybrid centromere region. A 800 bp region from pAWRC (AF245032) that had no similarity to CRWs was used to define a 9.9 Mb region on the 1RS side of the 1RS.1BL centromere while CRW/CCS1 (AB048244.1, a 249 bp repetitive unit), and Tail 1 (AB016967) from the wheat centromere were used to define the 1BL side of the 1RS.1BL chromosome (Figure 2a) (Francki, 2001; Keeble-Gagnère et al., 2018). In total the region covered by these centromere markers was 10.87 Mb within a region of 272.02 Mb to 296.20 Mb (24.18 Mb). The centromere marker sequences are evident in the matrix analysis (Figure 2a) as large arrays of repetitive sequences. At the junction between 1RS and 1BL, to form the 1RS.1BL chromosome, there exists a sharp change-over from the blocks of repetitive sequences carrying pAWRC (Figure 2a-a1, blue dashed line box) to the CRW markers sequences (Figure 2a-a1, red dashed line boxes). The matrix defines the junction between 281.93 Mb and 281.99 Mb and is between a RLG_Taes_Abia_B_3Brph7-445 element (coordinates 281,918,330 to 281,927,173 bp) and a LTR, Gypsy; consensus sequence (coordinates 281,926,633 to 281,935,207 bp) using TREP database (http://botserv2.uzh.ch/kelldata/trep-db/) to identify the LTR elements. The junction can be most easily modeled as resulting from a recombination event involving an Abia sequence located in Abia-like segments within CRW/Cereba elements in the wheat centromere and an Abia element in the original 1R chromosome.

**Figure 2.**
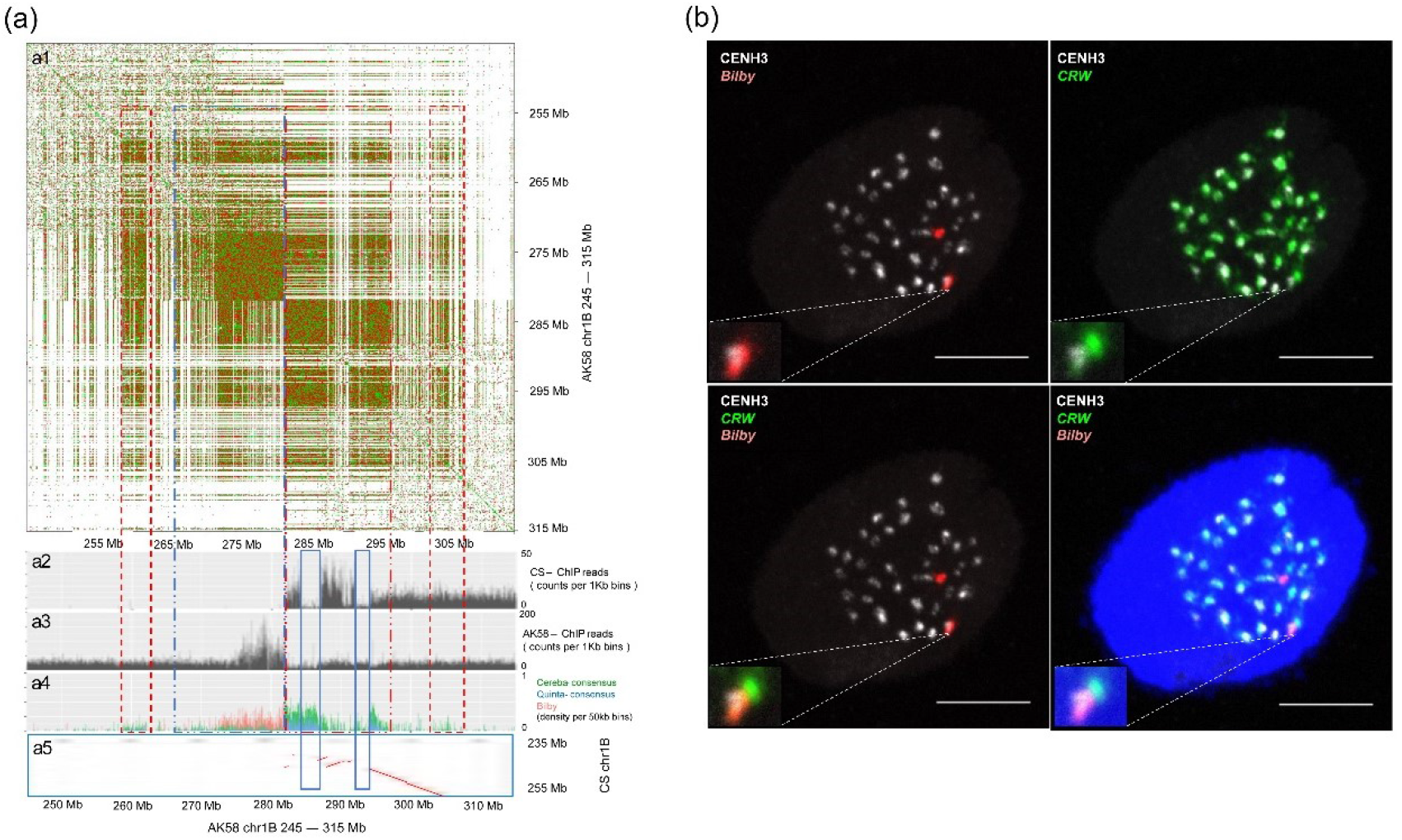
The 1RS.1BL centromere. (a) Dot-plot analysis of genome sequence at 245 Mb-315 Mb and BLAST-based distribution of ChIP-Seq data based on the CENH3-precipitation in AK58 and CS. (a1) Dot matrix of the AK58 centromere region of 70 Mb using YASS with the dashed-line boxes indicating the very large blocks of repetitive sequences (Kucherov et al., 2006). (a2) Blast of CENH3-antibody ChIP-Seq reads from CS on the 1RS.1BL centromere region showed two sub-domains which basically consist of the *CRW* and *Quinta* TEs. (a3) Blast of CENH3-antibody ChIP-Seq reads from AK58, the blast domain appears on the 1RS side (red) instead of 1BL relative to the wheat CRW (green) and Quinta (blue). (a4) Distribution of retrotransposons in the 1RS.1BL centromere region of AK58 - the boxes in dashed lines and solid lines are discussed in the text. (a5) Dot matrix of the 70 Mb AK58 centromere region vs the 20 Mb core centromere region from CS. The solid blue line boxes define regions addressed in the text. (b) Late prophase nuclei (same in each frame) show the *in situ* co-localization of the CENH3 antibody (white), the rye Bilby sequence (red) and the Cereba (CRW, green) sequences. The Bilby sequence (red) detects 1RS centromeres from the other 40 wheat centromeres (green). The green dots conjugating with the red dot is the 1BL centromere. The CENH3 signals were mainly co-located with the Bilby signals and this directly supports the centromere shift in observed in Figure 2 (a3) in the BLAST analysis of AK58-CENH3 ChIP-Seq reads.

The ca 150 Mb region that is relatively poor in annotated gene models (coordinates 165 to 313 Mb, Figure 1a and Figure 2a, dashed line boxes) houses predicted genes that code for peptides less than 50 amino acids and has a good coverage of hits from RNA-seq data originating from a range of tissues that could not be clearly assigned to gene models. Immunoprecipitation of CENH3 binding genome sequences from CS and AK58 nuclei provided a class of sequence to further define the core 31 Mb centromere region more clearly. The CS-CENH3 sequences differentiated the wheat centromere segment on 1BL from 1RS when aligned across the AK58_chr1RS.1BL_v6 assembly (Figure 2a-a2, coordinates 282 to 292 Mb) and also identified two sections of non-centromere (wheat) DNA (blue solid line boxes in Figure 2a-a5) even though the well-known centromere transposable elements, Cereba and Quinta, exist in these regions (Figure 2a-a1 and Figure 2a-a4). The AK58-CENH3 sequences mainly identified the rye centromere segment 272 to 282 Mb on 1RS (Figure 2a-a3). The dot matrix of CS vs AK58 core centromere sequences (Figure 2a-a5) indicated a region where the two genomes are structurally rearranged relative to each other. The *in situ* localization of the centromere Abia (rye) and CRW (wheat) sequences as well as the CENH3 protein using fluorescent antibodies (Figure 2b) indicated that the AK58 CENH3 sequences mainly co-located with the Bilby sequences (rye, Abia TE family) and confirmed that the CENH3-ChIP protocols selected specific sub-populations centromere sequences.

### Expression of 1RS genes in a wheat background

Transferring alien chromosome or chromosomal fragments from wheat relatives to wheat is an efficient approach for wheat improvement, relying on the expression of the alien chromosome genes in a wheat background. There are 1,480 high confidence genes annotated in 1RS and 1,560 on 1BS in CS (tissue expression summarized in Figure 3a) and entries of particular interest relate to the deployment of the 1RS.1BL chromosome in wheat breeding with a focus on those affecting grain quality and yield gains. Our study using the RNAseq data sets indicated that the 1RS gene expression is generally successful in wheat backgrounds and that 1RS genes do not interfere with wheat gene expression in total. A graphical representation of tissue-based expression groups is summarized in Figure 3a and 3b for AK58 and CS. At this broad level it is evident that 1RS has a greater percentage of genes expressed specifically in the grain tissue relative to CS (Figure 3b). In Figure 3c, the tissue specific gene expression patterns among the AK58 and CS chromosome 1 pairs are compared in more detail and it is evident that only 1BS in AK58 (= 1RS), has higher levels than 1BS of CS at the grain-20 DPA stage, where we found 6.44× higher levels of total gene expression in 1RS of AK58 relative to 1BS in CS. This grain-20 DPA stage is the active grain filling stage and the higher levels of gene expression may thus relate to the large grain size (45 g/1000 grain) and higher yield (7.5 t/h) of AK58 compared to the small grain size (35 g/1000 grain) and lower yield (3.0 t/h) of CS.

**Figure 3.**
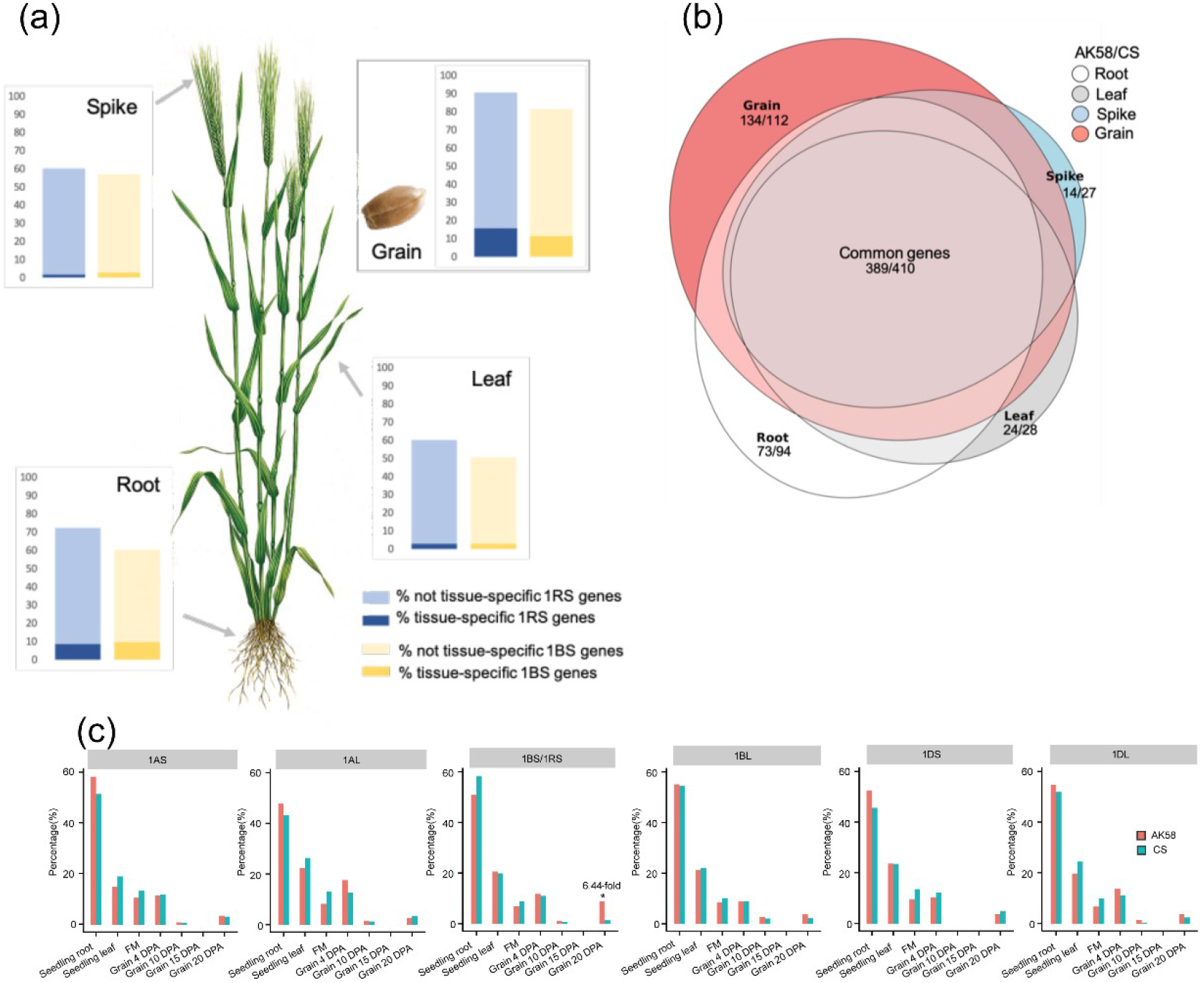
Comparative analysis of gene expression in group 1 chromosomes between AK58 and CS. (a) Expression pattern of genes in 1RS (blue) and 1BS (yellow) shown as % of total gene number expressed in the spike, root, grain and leaf tissues. Tissue-specific genes are highlighted in darker shades both for 1RS (blue) and 1BS (yellow). (b) Comparison of tissue-specificity in AK58 and CS with the numbers showing information for each line and tissue as AK58/CS. (c) Comparison of tissue specific gene expression across the short and long arms of the group 1 chromosomes. The 1BS/1RS panel highlights a major difference between AK58 and CS in the grain-DPA20 stage where a 6.44-fold higher expression of transcripts in AK58 vs CS at 20DPA was identified.

The hierarchical clustering of gene expression in Figure S4a within the AK58-1RS gene space of the developing grain provides a greater resolution the view of the gene families during the progressive differentiation of the grain tissue occurs as the spike matures and was consistent with IWGSC (IWGSC et al., 2018) and Ramírez-González (Ramírez-González et al., 2018). In Figure S4b, the early, prominent, expression of the MIKC-type MADS-box transcription factor family was striking. Genes that were highly expressed in grain tissue were of interest because historically the presence of 1RS in bread wheat was considered detrimental for qualities relating to the performance of flour from these wheat lines in the standard processing methodologies (Lukaszewski et al., 2000; Gobaa et al., 2007; Li et al., 2016). Transcription factor gene families prominently expressed in roots (as well as leaves) such as Cys3His zinc finger protein (C3H), MYB protein family (MYB) and WRKY signature containing transcription factor (WRKY) (Figure S4b) were of interest because this class of gene was well represented in the region that was studied by Howell et al. in which disruption of the genome structure through recombination with wheat 1BS reduced yield (more detailed analysis below) (Howell et al., 2019).

### The pentatricopeptide-repeat (PPR) gene family at the *rf/Rf ^multi^* loci of 1RS and 1BS

PPR domain carrying proteins form the one of largest gene families of land plants and are involved in the regulation of RNA metabolism including RNA editing, stability, processing, and splicing to translation (Lurin et al., 2004; Cheng et al., 2016). Most of restorer of fertility (*Rf*) genes cloned in model organisms encode P-class PPR proteins. AK58 is a 1RS.1BL variety and the 1RS.1BL translocation replaces *Rf ^multi^* locus from chromosome 1BS of wheat (15.28 to 59.84 Mb in CS) in multiple CMS systems (*Aegilops kotschyi, Ae. uniaristata* and *Ae. mutica*) to generate male sterile wheats (Tables S3 and 4) (Lukaszewski et al., 2017; Tsunewaki, 2015). The syntenic region in AK58-1RS is at 0.58 to 52.35 Mb in our assembly (Tables S5 and 6). The overall gene numbers in the regions that satisfy the definition of the PPR gene models are 21 in CS and 8 in AK58(Tables S3-5). A cluster of 11 PPR-family genes in CS-*Rf ^multi^* are a tandem array (at location 56.49 to 58.34 Mb) and a similar cluster exists in AK58-*rf ^multi^* (at 48.60 to 49.42 Mb) but is not in an exactly matching syntenic location (Figure 4a, Tables S6 and 7). One gene model, *TraesCS1B01G072900,* from the CS cluster has a homologous sequence *TraesAK58CH1B01G045250* at 49.41 Mb in AK58 and this gene in AK58 is also syntenic to *TraesCS1B01G074600* in CS-*Rf ^multi^.* The *TraesCS1B01G074600* gene is, in turn, syntenic to another AK58 gene (*TraesAK58CH1B01G043900*) in the AK58-*rf ^multi^* region, so we consider it useful to define this as one closely related group represented by *TraesCS1B01G072900*. These relationships reflect a complex micro-level relationship between the AK58-*rf ^multi^* and *CS-Rf ^multi^* regions which results in the different expression patterns that are relevant to the male fertility/restoration of fertility (Figure 4b). The three P-class proteins from the *CS-Rf ^multi^* region, namely, *TraesCS1B01G072300, TraesCS1B01G072900* and *TraesCS1B01G074600,* and their 1RS homologs, have significant mitochondrial targeting scores (Figure 4c, Tables S6 and 8) and between these genes only *TraesCS1B01G072300* shows the most striking difference in transcription in FM tissue when compared to its homolog in AK58 (*TraesAK58CH1B01G045100*) (Figure 4b).

**Figure 4.**
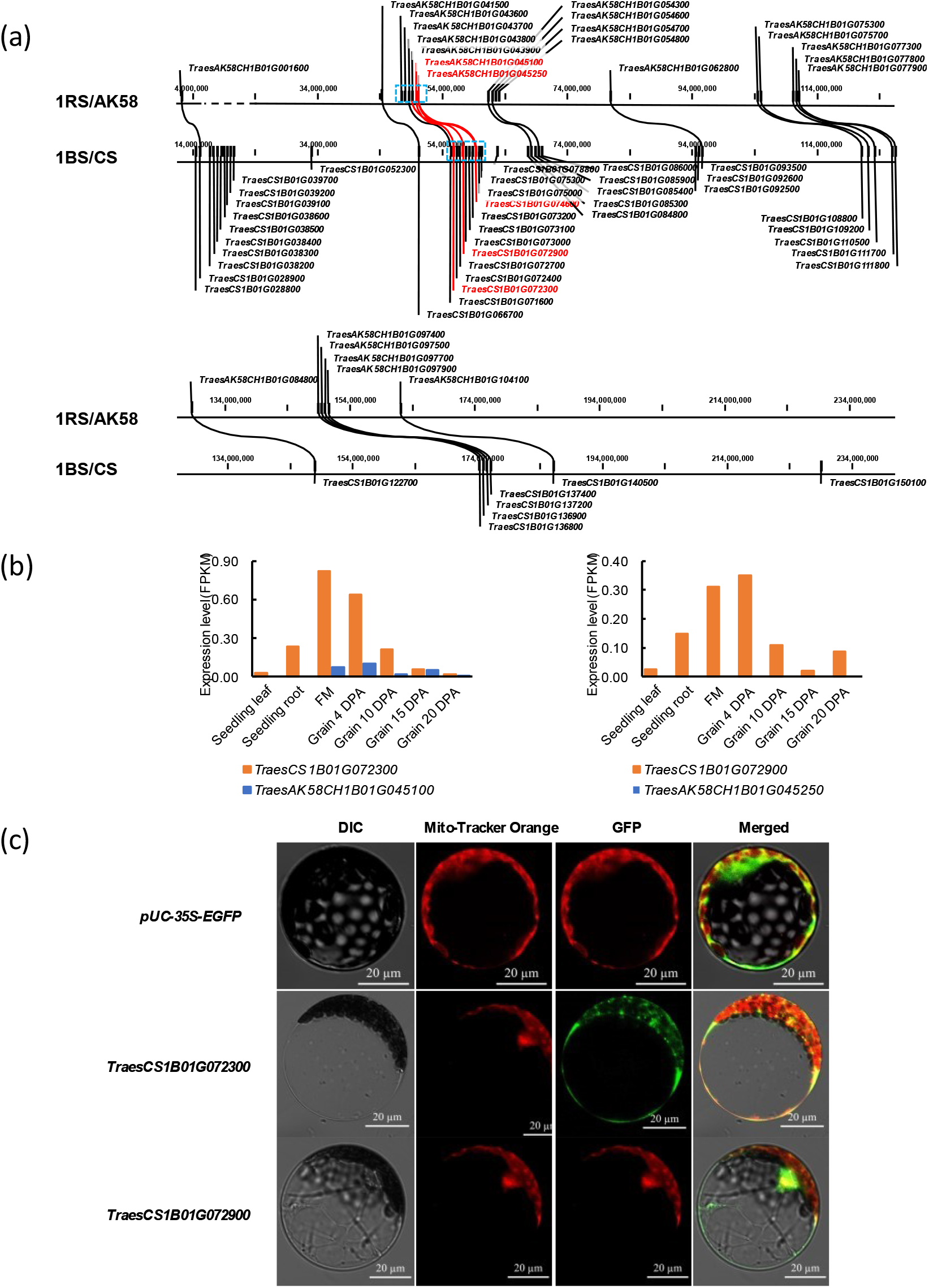
PPR gene comparisons between 1RS of AK58 and 1BS of CS. (a) Gene collinearity analysis. (b) PPR gene expression comparison for the genes indicated in either orange (CS origin) or AK58 (blue origin) for the tissues, seedling leaf, seedling root, FM (flowering meristem) and grain tissue (DPA = days post anthesis). (c) Subcellular localization of *TraesCS1B01G072300* and *TraesCS1B01G072900.* Protoplasts transformed with 35S: GFP (pUC-35S-GFP), 35S:TraesCS1B01G072300-GFP (*TraesCS1B01G072300*) and 35S:TraesCS1B01G072900-GFP (*TraesCS1B01G072900*) constructs were analyzed using fluorescence microscopy. DIC indicated bright field. The dye Mito-Tracker Orange was used as a mitochondrial marker. (Scale bars: 20 μm.).

Taken together, the *CS-Rf ^multi^* /AK58-*rf ^multi^* comparison suggests the *TraesCS1B01G072300* and *TraesCS1B01G072900* are the most likely candidate genes of the multi fertility restoring locus *Rf ^multi^* in the context of the 1RS.1BL translocation replacing the *Rf ^multi^* locus from chromosome 1BS in multiple *Aegilops* CMS systems that generate male sterile wheats and male fertility being subsequently restored by the 1BS-*Rf ^multi^* locus (Lukaszewski et al., 2017; Tsunewaki, 2015).

### The grain storage protein gene families

Gamma and omega secalin sequences were identified and aligned to gamma and omega gliadins from the CS chromosome 1B. While the sequence similarity between gamma gliadins and gamma secalins was over 70%, omega gliadin and omega secalin sequences differed significantly both in the signal peptide as well as in their repetitive regions. Nineteen gamma secalin coding sequences were identified from which 18 were located in a single cluster at 1RS between positions 8,722,719 bp and 8,904,808 bp (Figure 5a). Using an extensive proteome database of rye grain peptides (Figure S1 and Methods) (Bose et al., 2019), half of these genes were verified at peptide level as bone-fide gamma secalin protein coding genes. Eight sequences (7 gamma secalins, 1 purinin) represented complete sequences, two sequences were partial sequences and ten sequences contained frameshifts or internal stop codons. Similarly, a cluster of 18 omega secalin genes were identified, between positions 18,457,234 bp and 18,690,191 bp with 13 verified at the proteome level (Figure 5b). Between the gamma secalin and omega secalin loci a single purinin gene was identified in a conserved position compared to chromosome 1B.

**Figure 5.**
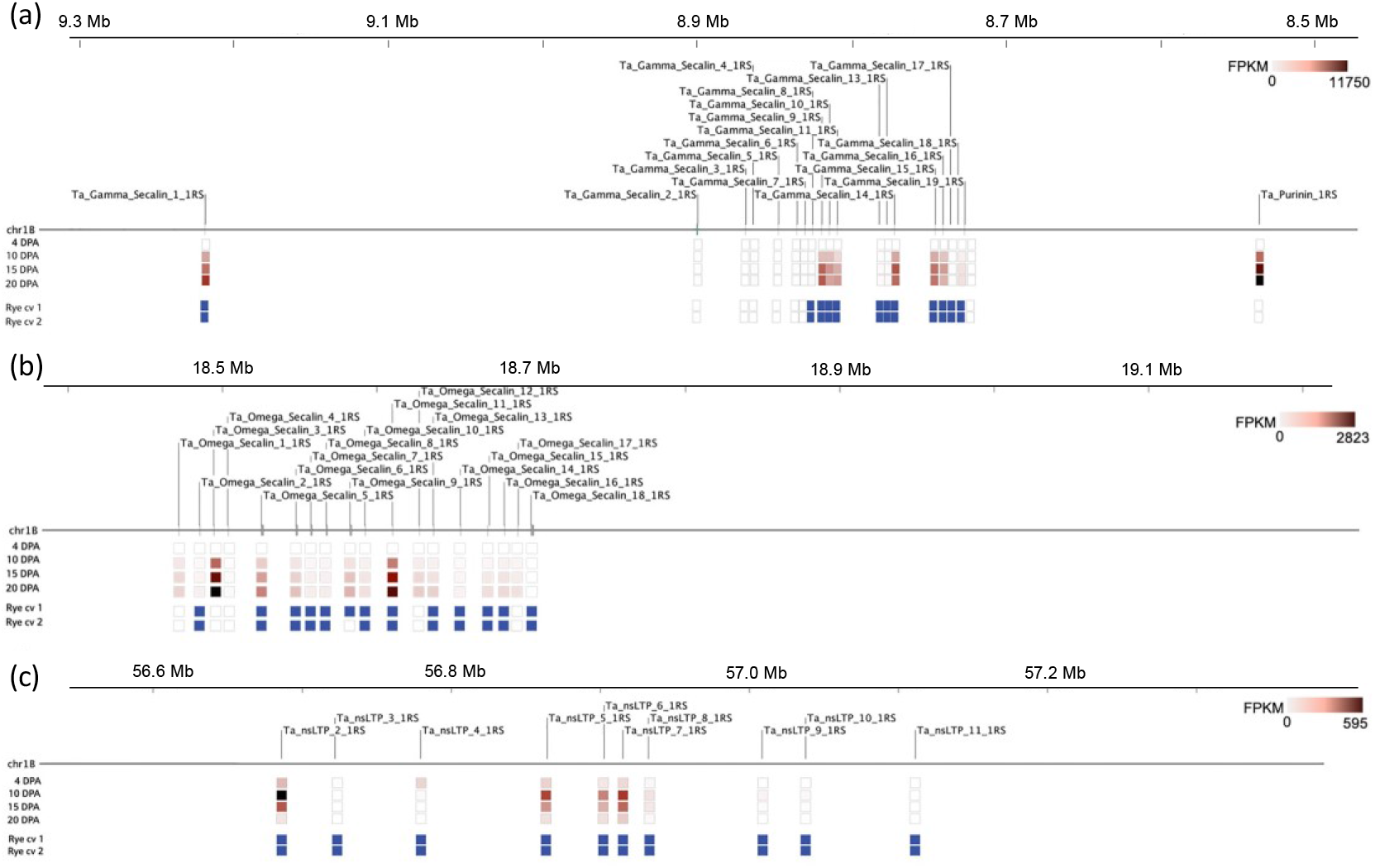
The grain storage protein gene analysis. (a) Gamma secalin locus on chromosome arm 1RS. (b) Omega secalin locus on chromosome arm 1RS. (c) nsLTP locus on chromosome arm 1RS.The tracks underneath each of genome map locations provide the level of expression at the respective stages of grain development and evidence for the representation of the gene models as grain proteins in extensive rye proteome studies.

Within the identified gamma secalins there were two sequences (Gamma secalin 5 and Gamma secalin 16) with Tryp_alpha_amyl domain (PF00234) while the majority of gamma secalins possess a single Gliadin Pfam domain (PF13016). In bread wheat, all the identified gamma gliadins and low molecular weight (LMW) glutenins have PF13016 (Gliadin) domains, and Tryp_alpha_amyl domains were only found in alpha gliadins among the gliadin and glutenin types. The Pfam domain composition of available secalin sequences from UniProt DB indicates the Gliadin domain (PF13016) and Tryp_alpha_amyl domain (PF00234) are both characteristic of rye secalins, while PF00234 domains were only identified in the 75 k gamma secalins. The Tryp_alpha_amyl domain containing proteins represent a sulphur-rich domain structure and are primarily characterized to have functions related to defense mechanisms against pathogens or lipid transport (non-specific lipid transfer proteins). As both in 1BS and 1RS the major storage proteins are located within regions enriched in disease resistance proteins this might indicate a potential alternative function of these proteins and their possible involvement in stress responses.

There were eleven nsLTP genes found on the 1RS chromosome arm, ten of which represented the PR60 nsLTP sub-type (UniRef100_B2C4K0) clustered between 56.6 Mb and 57.2 Mb and specifically expressed in grain tissue (Figure 5c); a single nsLTP (at 42.24 Mb) was highly expressed in roots. On the long arm, 7 nsLTPs, 5 LTPs and 4 LTP-like sequences were identified from which 3 short LTP-like sequences were grain specific. All the prolamin genes present on 1RS were specifically/preferentially expressed in the grain, while the nsLTPs were also expressed in other tissues such as under-spike internodes and young spikes.

### The disease resistance gene families

The *Pm8* gene, orthologue of the *Pm3,* provided a good model for the RGA gene in the region although the expression of the gene itself was not confined to any single tissue category (Hurni et al., 2013). The identification of *Pm8* suggested that 1RS of AK58 is from diploid rye Petkus. The RGA genes specifically expressed in different tissue were assessed against a total of 2,871 gene models with the NB-ARC domain in their structure were identified within the IWGSC RefSeq v1.0. In the AK58 1RS.1BL chromosome, disease resistance gene models were identified (Table 1, Figure S1 and Methods). The NB-ARC domains are the most common feature within disease resistance genes and are key components of apoptosomes that are involved in recognizing the presence of a pathogen or DNA damage in the cell and responding to the problem by localized cell death to protect the organism per se (Crespo-Herrera et al., 2017; Li et al., 2016). The range of gene diversity is consistent with the range of intrinsic and extrinsic stimuli to which wheat is exposed. Among the 22 1RS-specific gene models we identified three broad categories, dominated by a large group (15) which comprised members of RGA families located on wheat 1B as well as other chromosomes and included genes that were identified as having disease resistance-like protein and tyrosine kinase domains (Table S9). Four gene models were part of a very large family encoding proteins with lrr-serine/threonine kinase domains. Three gene models were of particular interest in that they were absent from 1BS and thus represent novel resistance genes introduced by 1RS (details in Table S9). Two of these gene models *TraesAK58CH1B01G008800* (at 4,676,665 bp) and *TraesAK58CH1B01G010100* (at 6,010,958 bp) are closely linked to, and on the proximal side of, the marker BE405749.1 which defines a region housing *Lr26* (Mago et al., 2002), as well as having the protein fold c6j5tc associated with the disease resistance gene rpp13-like protein 4, determined using Phyre2 (Kelley et al., 2015).

**Table 1.**
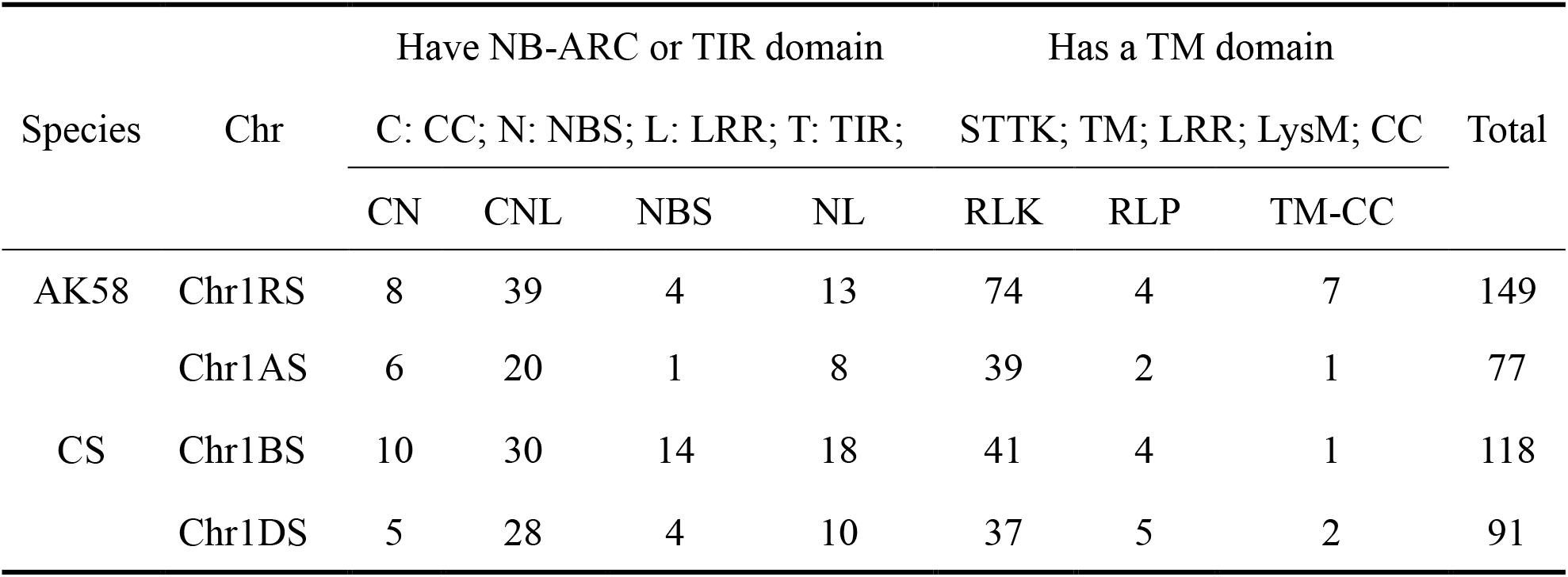
RGAs prediction in AK58 1RS compared to CS 1AS, 1BS and 1DS, respectively.

### Characterization of the 1RS region disrupted by recombination between 1RS and 1BS

The yield-related region (YR) comprised the genome region in the terminal 14 Mb on the 1RS.1BL chromosome based on published molecular marker evidence in Mago et al. and Howell et al. (Mago et al., 2002; Howell et al., 2014). The region exists within the terminal 22 Mb and housed 259 genes, from a total of 465, that could be characterized by the clusters of co-expression representing gene networks potentially disrupted by recombination events between 1RS and 1BS. The analysis in Figure 6 shows the co-expression matrix of the contig-356, −445, −624 and −517 1RS genes (Table S10). The gene expression was calculated using the Morpheus analysis tool (https://software.broadinstitute.org/morpheus/) and a 0.5 FPKM cut-off. The co-expression similarity matrix was calculated using Pearson correlation and the clustering was carried out using only the co-expression values > absolute value 0.7 FPKM as being significant (highlighted red = positively correlated, blue = negatively correlated). The contigs annotated as “Tissues” were identified based on the stage and tissue-specific clustering in Figure 6. The boxed regions identified gene clusters that formed qualitatively major networks based on shared patterns of expression. The 1RS genes in clusters 1-7 shown in Figure 6 are combinations of models that are tissue specific in expression and ones that are more generally expressed and overall, the analysis indicates that the genes in the YR region are widely networked to genes involved in a broad range of biological activity with a particular focus on root specific genes (Table S10). The clusters also include genes potentially interacting with nitrogen metabolism related genes as well as genes involved in stress responses and adaptation and thus a disruption in the activity profiles is predicted to have wide ranging effects on a complex phenotype such yield. In particular we identified TraesAK58CH1B01G010700 (formin-like domain protein) and TraesAK58CH1B01G007500 (Cathepsin-L, homolog) genes as having the highest number of co-expression partners. The rice homolog for the *TraesAK58CH1B01G010700* gene is *Os05t0104000-00,* a formin-like-domain protein FH14, and defines a class of protein which interacts with microtubules and microfilaments to regulate cell division. This gene also shows co-expression with a nitrate reductase at chromosome 6B. The *TraesAK58CH1B01G010700* gene showed a broad co-expression-based network including interactions with auxin response factors on chromosomes 1A, 1B, 1D, 2B, 5A, 7A, 7B, 7D.

**Figure 6.**
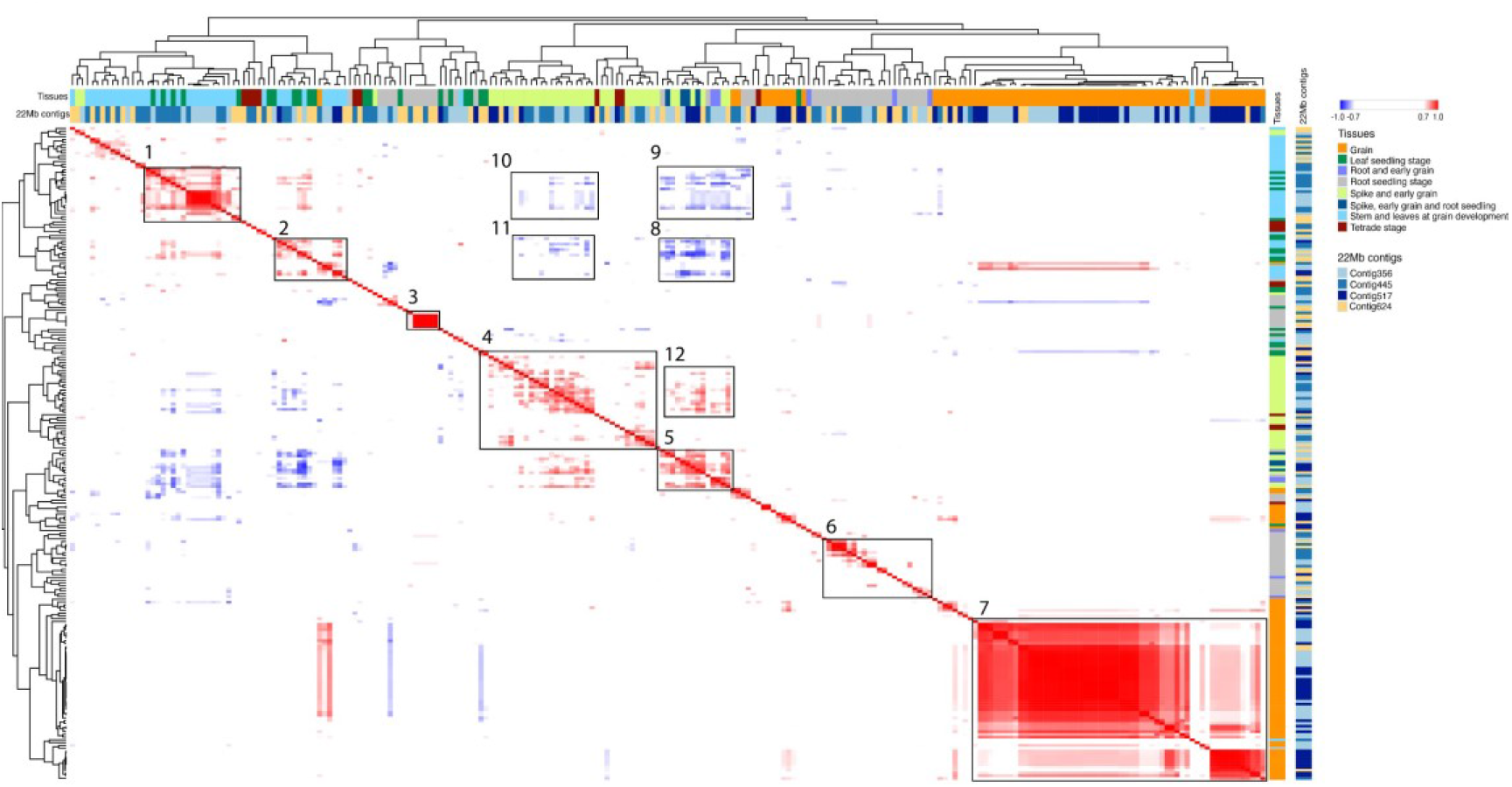
Co-expression analysis of homologous genes expressed in the 22 Mb region of AK58 1RS. Co-expression similarity values are represented in AK58 1RS calculated using Pearson correlation followed by hierarchical clustering using the pre-computed similarity matrix values. Co-expression values below −0.7 are labelled in blue and represent strong negative co-expression. Co-expression above 0.7 represents strong expression similarity and is highlighted in red. Tissue specificity and location of the genes within the analyzed contigs in the 22 Mb region are highlighted with different colors. Matrix was generated using https://software.broadinstitute.org/morpheus/.

## DISCUSSION

Our study provides the genome assembly for chromosome 1RS.1BL, a chromosome which was one of the early successes of genetic engineering at a chromosome level that has had a significant impact on wheat yield and hence food production globally (Schlegel and Korzun, 1997). The value of our assembly is demonstrated through the identification of 1RS-specific genes within an overall gene space that showed excellent synteny with 1BS from wheat. In contrast the non-gene space is shown to have regions of highly amplified, relatively short (100 – 500 bp), units of DNA sequences in regions which are not syntenic to equivalent regions in 1BS and are suggested to be possible sites for recombination type events between 1R and 1B. The detailed analysis of the centromere identified the junction between 1RS and 1BL and this was found to comprise part of an Abia transposable element (TE) on the 1RS side and a Cerebra TE on the 1BS side suggesting that a recombination event between the arrays of Abia TEs that characterize the rye centromere and the arrays of Cereba TEs in the 1B centromere, which can also carry Abia-like sequences (http://botserv2.uzh.ch/kelldata/trep-db/), could have generated the 1RS.1BL chromosome (Francki, 2001). Our analysis of the CENH3 locations indicated a shift in the location of the centromere assembly defined by CENH3 to the rye side of the translocation and provided a clear indication of mobility in the location of the point of attachment of micro-fibrils for mitosis.

Specific families of gene models characterized in detail in this study included the genes predicted to be involved in the male sterile/male fertility restoration interactions, resistance gene analogs and the gamma and omega secalin storage proteins. The substitution of 1BS with 1RS in 1RS.1BL wheat lines results in the replacement of three PPR proteins that have high mitochondria target scores. In the context of the *CS-Rf ^multi^* /AK58-*rf ^multi^* comparison carried out, it is suggested that *TraesCS1B01G072300* and *TraesCS1B01G072900* are representative of the most likely candidate genes of the multi fertility restoring *Rf ^multi^* locus because they show the most striking differences in transcription to its AK58 homolog and could thus be most significant in restoring male fertility the 1BS-*Rf ^multi^* locus in the multiple CMS systems-male sterile wheats where 1RS.1BL translocation replaces the *Rf ^multi^* locus from chromosome 1BS (Lukaszewski et al., 2017; Tsunewaki, 2015). Our analysis of the resistance gene analogs identified three genes that were absent from 1BS and thus represent new resistance genes for varieties carrying the 1RS.1BL translocation. Two of these genes, *TraesAK58CH1B01G008800* (at 4,676,665 bp) and *TraesAK58CH1B01T0G0100* (at 6,010,958 bp), locate in the genome region housing genetically defined disease resistance genes and are thus candidate genes for entities such as *Lr26*.

The substitution of 1BS in 1RS.1BL wheat lines also results in the replacement of a major source of wheat gliadin proteins on 1BS with secalin protein coding genes on the 1RS.1BL chromosome. The analysis of the gamma and omega secalin storage protein coding regions and expression identified pseudogenes as well as two sequences (Gamma secalin 5 and Gamma secalin 16) with Tryp_alpha_amyl domain (PF00234). The Tryp_alpha_amyl domain containing proteins, represent a sulphur-rich domain structure are more broadly associated with defense mechanisms against pathogens, lipid transport and storage function and suggests an involvement in stress responses. In the context of unwanted processing attributes (“sticky” dough) associated with flour from 1RS.1BL containing wheat cultivars genes related to the production of xylose and arabinoxylose and the xylanase domain protein (TraesAK58CH1B01G088700) specifically expressed at 4 DPA in the grain could be related to the moderation of arabinoxylans that may be associated with increased water absorption resulting in the “sticky” dough defect (Gobaa et al., 2007; Henry et al., 1989; Lee et al., 1995).

The mapping of published DNA probes to the 1RS assembly indicated that a 9 Mb region (in the terminal 14 Mb, “YR” region) was disrupted by the 1RS/1BS recombinants selected by Lukaszewski (Lukaszewski et al., 2000). The genes in this region are widely networked, as expected, based on co-expression analyses and have a striking representation of root-specific genes that are good candidates for relating the changes in root phenotype to the disruption of the 9Mb region defined by Howell et al (Howell et al., 2014 and 2019). Our study highlighted the complex negative and positive interactions among the genes in this YR region and identified TraesAK58CH1B01G010700 (formin-like domain protein) and TraesAK58CH1B01G007500 (Cathepsin-L, homolog) genes as having the highest number of co-expression partners. The rice homolog for the *TraesAK58CH1B01G010700* gene is *Os05t0104000-00,* a formin-like-domain protein FH14, showed a broad co-expression-based network across 8 chromosomes consistent with a gene model that encodes a protein regulating cell division.

In brief, we generated a high quality 1RS.1BL translocation chromosome sequence which provided a basis for defining gene underpinning the agronomic attributes of the 1RS.1BL translocation chromosome in wheat improvement. The structure, gene complement of 1RS.1BL and candidate genes identified in this study provides the resource-of-choice for refining the contribution of this chromosome to wheat genetic improvement.

## EXPERIMENTAL PROCEDURES

### Gene annotation

Protein-coding identification and gene prediction were carried out using a combination of homology-based prediction, *de novo* prediction, and transcriptome-based prediction methods (Figure S1). Five *ab initio* gene prediction programs, Augustus (v.2.5.5), Genscan (v.1.0), GlimmerHMM (v.3.0.1), Geneid, and SNAP, were used to predict coding regions in the repeat-masked genome. Proteins from ten plant genomes (*T. aestivum*, *T. turgidum dicoccides, T. urartu, Ae. tauschii, Hordeum vulgare, Brachypodium distachyon, Oryza. sativa, Zea mays, Sorghum bicolor* and *Setaria italic*) were downloaded from EnsemblPlants (http://plants.ensembl.org/index.html). *Panicum virgatum* genome was downloaded from Phytozome (https://phytozome.jgi.doe.gov/pz/portal.html). Protein sequences from these genomes were aligned to the AK58 assembly using TblastN with an *E-value* cutoff of 1e-5. The BLAST hits were conjoined using Solar software. GeneWise was used to predict the exact gene structure of the corresponding genomic regions for each BLAST hit. Homology predictions were split into two sets, which included a high-confidence homology set (HCH-set) with predictions from genomes with CS wheat and a low confidence homology set (LCH-set).

A collection of wheat FLcDNAs (16,807 sequences) were directly mapped to the AK58 genome and assembled by Program to Assemble Spliced Alignments (PASA). Gene models created by PASA were denoted as the PASA-FLC-set (PASA full length cDNA set), this gene set was used to train the *ab initio* gene prediction programs. RNA-seq data were mapped to the assembly using Tophat (v.2.0.8). Cufflinks (v.2.1.1) was then used to assemble the transcripts into gene models (Cufflinks-set). In addition, a total of 2,016 Gb RNA-seq data from different organs (root, leaf, internode, flower and developing seed) were assembled by Trinity, creating several pseudo-ESTs. These pseudo-ESTs were also mapped to the AK58 assembly and gene models were predicted using the PASA. This gene set was denoted as PASA-T-set (PASA Trinity set).

Gene model evidence from the HCH-set, LCH-set, PASA-FLC-set, Cufflinks-set, PASA-T-set and *ab initio* programs were combined by EvidenceModeler (EVM) into a non-redundant set of gene structures. Weights for each type of evidence were set as follows: HCH-set > PASA-FLC-set > PASA-T-set > Cufflinks-set > LCH-set > Augustus > GeneID = SNAP = GlimmerHMM = Genscan. Gene model output by EVM with low confidence scores was filtered using: (1) coding region lengths of 150 bp, and (2) supported only by *ab initio* methods and with FPKM<1.

In an approach similar to that described for *Gossypium raimondii* genome studies (Yan et al., 2016), we further filtered gene models based on Cscore (Cscore is a peptide BLASTP score ratio mutual best hits BLASTP score), peptide coverage (coverage is highest percentage of peptide aligned to the best of homologues) and overlap of its CDS with TEs. The Cscore and peptide coverage were calculated as described in *G. raimondii* (Yan et al., 2016). Only transcripts with a Cscore ≥ 0.5 and peptide coverage ≥ 0.5 were retained. For gene models with more than 20% of their CDS sharing an overlap with TEs, we required that its Cscore must be at least 0.8 and that its peptide coverage must be at least 80%. Finally, we also filtered out gene models of which more than 30% of the peptides in length could be annotated as Pfam or Interprot TE domains.

### Transcript analysis

Two transcriptome datasets derived from AK58 genome sequencing project were used for this analysis. (1) 42 diverse samples (each 3 biological replicates, total 126 libraries) were collected for AK58, covering anther development to the tetrad stages, floret/spikelet meristem, three stages of stem development, three stages for flag leaf, five stages of grain development, and 7 day seedling for the leaf and root sample under normal or six abiotic stresses conditions. (2) Seven tissues shared between AK58 and CS (each 3 biological replicates, total 42 libraries), covering 21 days seedling for the leaf and root, floret meristems stage, and four grain development stages (4/10/15/20 days after flowering).

To align RNA-seq sequences to the genome assembly, the BLAT software within Apollo was used in two modes (Lee et al., 2013), one allowing only perfect matches and a second mode using the Hi-sat-2 default settings (https://ccb.jhu.edu/software/hisat2/index.shtml) equating to approximately 80% similarity over at least 80% of the sequence. Only unique gene models were used to define the 1RS.1BL synteny.

Gene co-expression matrix of the contig-356, −445, −624 and −517 1RS genes were developed using the Morpheus matrix visualization and analysis tool (https://software.broadinstitute.org/morpheus/). Mean values of gene expression data obtained from the different tissue samples with a 0.5 FPKM cut-off were used to calculate Pearson correlation coefficients. The obtained similarity matrix was used for hierarchical clustering with complete linkage. Rows and columns represent the individual gene models present in the analyses 22 Mb region (Table S10). Tissue specific expression patterns and location within the analyzed region are annotated by different colors were obtained from the analysis presented in Figure 6. Co-expression similarity values > absolute value 0.7 FPKM as being significant are highlighted in red (positively correlated) and blue (negatively correlated).

### Chromosome preparation

The protocol for root tip mitotic metaphase chromosomes of AK58 was largely referred to the previously report (Han et al., 2006). Briefly, the roots of AK58 grew to 1.5-2.0 cm in length were excised and treated with nitrous oxide gas for 2 h under pressure under 1 MPa. The treated roots were fixed in ice-cold 90% acetic acid for 10 min. Subsequently, the root tips were dissected and digested by 2% cellulose Onozuka R-10 (Yakult Pharmaceutical, Japan) and 1% pectolyase Y23 (Yakult Pharmaceutical) solution for 45 min at 37°C. After digestion, the root sections were broken in a 90% acetic acid. The cell suspension was dropped and air-dried on glass slides for chromosome observation.

### Fluorescence *in situ* hybridization

The method of sequential FISH and Non-denaturing FISH (ND-FISH) with different labeled probes for karyotype analysis of AK58 was mainly performed according to the previously published protocol (Fu et al., 2015). The probe Sec1 for rye specific secalin was labeled with Texas Red-5-dUTP (Invitrogen) or Alexa Fluor 488-5-dUTP (Invitrogen) using nick translation for FISH (Clarke et al., 1996). The oligonucleotide probes for ND-FISH with centromeric specific probe CCS1, 18S-45SrDNA probe pTa71, and probe pSc119.2 were referred to Tang et al. (Tang et al., 2014). The repeats probes pSc200 and 5SrDNA were from the previously published information (Fu et al., 2015; Lang et al., 2019). The synthetic oligo probes were 5’ end-labeled with 6-carboxyfluorescein (FAM) for the green signal and 6-carboxytetramethylrhodamine (Tamra) for the red signal. The slides after FISH and ND-FISH were mounted with Vectashield mounting medium containing 1.5 μg/mL 4, 6 - diamidino −2- phenylindole (DAPI, Vector Laboratories, Burlingame, CA, USA). The FISH images were captured with an Olympus BX-53 microscope equipped with a DP-80 CCD camera.

### 660K SNP Analysis of the 1RS Lines and Non-1RS Lines

Wheat 660K SNP Array designed by the Institute of Crop Sciences of the Chinese Academy of Agricultural Sciences and synthesized by Affymetrix® was applied to genotype 36 1RS.1BL lines and 9 non 1RS.1BL lines. All the SNPs were merged to FASTA file for 45 samples. TreeBeST (1.9.2) (http://treesoft.sourceforge.net/treebest.shtml#intro) nj was used to build neighbor-joining phylogenetic tree with parameters: “-b 1000” and MEGA7 was used to visualize phylogenetic trees (Kumar et al., 2016).

### ChIP-seq

We used chromatin immunoprecipitation and sequencing technique to find the centromeric DNA by CENH3 antibody, which was a rabbit polyclonal antiserum and was raised against the peptide ‘CARTKHPAVRKTK’ (Li et al., 2013). ChIP was conducted using young leaves of AK58 as previously described (Nagaki et al., 2003). The enriched DNA samples were sequenced using Illumina Hiseq X-10 to generate 150 bp paired-end sequences. Reads were filtered with TrimGalore (http://www.bioinformatics.babraham.ac.uk/projects/trim_galore/) and aligned to the AK58 genome sequence using Bowtie 2 (Langmead and Salzberg, 2012). We only retained reads that determined the unique position, which MAPQ >= 30, for further analysis. The distribution of ChIP-seq reads were calculated using the unique read number per 1 kb window. ChIP-Seq data precipitated from CS in our previous study (SAMN11655702) were analyzed as parallel.

### Centromeric sequence analysis

In order to detect the sequence variation, we aligned the sequence of 1B from CS to AK58 genome using NUCmer program (parameter: -c 700) in MUMmer4 package (Marcais et al., 2018). We used mummerplot program in the same software to draw 245–315 Mb of 1B chromosome from AK58. Consensus of *CRW, Quinta* and *Bilby* were aligned to AK58 genome and calculated the percentage per 50 kb to reveal the sequence composition.

### Immuno-co-localization Analysis of CENH3 with *CRW* and *Bilby*

Root tips of young seedling were collected for Immuno-staining as previously described (Zhao et al., 2019). *CRW* and *Bilby* were labeled with biotin-16-Dutp and digoxigenin-11-dUTP via nick translation respectively. The hybridized probes were detected by fluorescein-conjugated goat anti-biotin and anti-digoxigenin-rhodamine Fab fragments coupled with TAMRA respectively. CENH3 was identified with anti-CENH3 antibody detected by Goat anti-Rabbit IgG (H+L) conjugated Alexa Fluor 647. Images were taken by a confocal (ZEISS LSM880) and processed using Adobe Photoshop CS.

### PPR gene analysis

The genome sequence data and gene annotation of CS were downloaded from EnsemblPlants (http://plants.ensembl.org/index.html). PPR sequences were annotated by Pfam databases (http://pfam.xfam.org/, v.32.0), and IPR database (https://www.ebi.ac.uk/interpro/, v.77.0), respectively. The classification of PPR was referred to Cheng et al. (Cheng et al., 2016). The PPR gene collinearity between 1RS of AK58 and 1BS of CS was analyzed by best reciprocal hit BLAST (*E-value* cutoff of 1e-5) (Gabriel and Kristen, 2007). The PPR gene’s subcellular localization was predicted using PSI (http://bis.zju.edu.cn/psi/) and validated by experiment. In briefly, full-length PPR cDNA was amplified and inserted in front of a GFP-coding sequence on the pUC-35S-EGFP vector. Tobacco (*Nicotiana benthamiana*) protoplasts were prepared and transfected according to the method described by Shan et al (2014). The protoplasts were subsequently stained with 10μm MitoTracker™ Orange CMTMRos (Invitrogen, M7510) for 10 min, and then examined on using a confocal scanning microscope system (ZEISS LSM880).

### The grain storage protein gene analysis

Prolamin super-family genes identified in the wheat reference genome were used to manually annotate the prolamin genes in the AK58 genome sequence (IWGSC et al., 2018; Juhász et al., 2018). Translated sequences were checked for the presence of signal peptides and the conserved cysteine pattern and Pfam domains as previously described (Juhász et al., 2018). Obtained sequences were aligned with gliadins and secalins retrieved from the Uniprot database to confirm the protein sub-types. Expression of genes was analyzed using the grain specific transcriptome data set obtained from 4, 10, 15 and 20 DPA grain libraries. Protein level expression of the translated secalins and nsLTPs were analyzed using the published data (Bose et al., 2019). LC-MS-MS data originally generated from tryptic digests of rye flour protein extracts were re-analyzed and protein identification was undertaken using ProteinPilotTM 5.0 software (SCIEX) with the Paragon and ProGroup algorithms with searches conducted against the Poaceae subset of the Uniprot database appended with the identified 1RS gene models and a contaminant database (Common Repository of Adventitious Proteins) (Shilov et al., 2007). Obtained fully tryptic peptides were mapped to the secalin and nsLTP sequences in CLC Genomics Workbench v12 (Qiagen, Aarhus, Denmark) using 100% sequence identity to confirm the expression at individual protein levels.

### RGA gene analysis

To predict RGAs in AK58 genome, a new plant RGAs database was constructed using protein sequences from the RGAdb (https://bitbucket.org/yaanlpc/rgaugury/src/master/) and candidate RGAs predicted in *Ae. tauschii* genome (Li et al., 2016; Luo et al., 2017). A total of 61372 disease resistance related sequences were obtained. Protein sequences of all annotated genes of AK58 were aligned to the new RGAs database using BLASTP with *E-value* cutoff of 1e-5. Potential RGAs were selected based on rule 80:80:80 (query sequence coverage more than 80%, target sequence coverage more than 80 and identity more than 80%). Eight RGAs-related domains and motifs including NB-ARC, NBS, LRR, TM, STTK, LysM, CC and TIR were searched and identified by RGAugury pipeline (Li et al., 2016). RGA candidates were predicted in *T. aestivum* (CS) genome using the same method.

## ACKNOWLEDGES

The authors are grateful to Yaoguang Liu and his laboratory team from the College of Life Sciences of the South China Agricultural University for suggestions in analyzing PPR gene family. This research was supported by The National Key Research and Development Program of China (2016YFD0101802 and 2016YFD0101602), Talent Program and Innovation Program of CAAS.

## AUTHOR CONTRIBUTIONS

ZR, DC, JJ, RA, and XK initiated the project and designed the study. AJ, DL, PD, ZJ, LF, KW, GKG, ZY, GL, DW, UB, MC, CK, GZ, XZ, XL, GC, YW, ZN, and LW performed the research. AJ, DL, PD, ZJ, LF, KW, GKG, CK, and GZ generated and analyzed the data. RA, XK, JJ, AJ, DL, PD, ZJ and KW wrote the paper.

## DATA AVAILABILITY

The whole genome sequence data reported in this paper have been deposited in the Genome Warehouse in National Genomics Data Center, Beijing Institute of Genomics (China National Center for Bioinformation), Chinese Academy of Sciences, under accession number GWHANRF00000000 that is publicly accessible at https://bigd.big.ac.cn/gwh. And the Assembly and availability of 1RS.1BL genome sequence can be found in URGI, https://urgi.versailles.inra.fr/download/iwgsc/AK58_1RS.1BL/.

## SUPPORTING INFORMATION

**Figure S1.** Protein-coding gene prediction process overview.

**Figure S2.** Neighbor-joining phylogenetic tree for SNP analysis of 36 1RS.1BL lines.

**Figure S3.** Distribution of some of the dominant retrotransposable elements in 1RS. The elements are identified on the right-hand side together with a color scale indicating the relative prominence of the elements.

**Figure S4.** Transcript analysis.

**Table S1.** Pedigrees of wheat lines used for 660k SNP analysis.

**Table S2.** 1RS dominant TE information.

**Table S3.** PPR genes identified on the CS genome and their expression levels in different tissues.

**Table S4.** PPR genes located on the region of *Rf ^multi^.*

**Table S5.** PPR Genes identified on the AK58 genome and their expression levels in different tissues.

**Table S6.** Orthologous PPR genes from 1BS (CS) and 1RS (AK58).

**Table S7.** Orthologous genes from 1BS (CS) and 1RS (AK58) on the region of *Rf ^multi^/rf ^multi^.*

**Table S8.** PCR primers used in this study for subcellular localization.

**Table S9.** RGA Gene models analysis on the AK58 genome.

**Table S10.** A summary of the gene models in the morpheus clusters for contig-356, −445, −517 and −624 in the 1RS.1BL assembly.

