## Supplemental FigureS1-4 for "1RS.1BL molecular resolution provides novel contributions to wheat improvement"

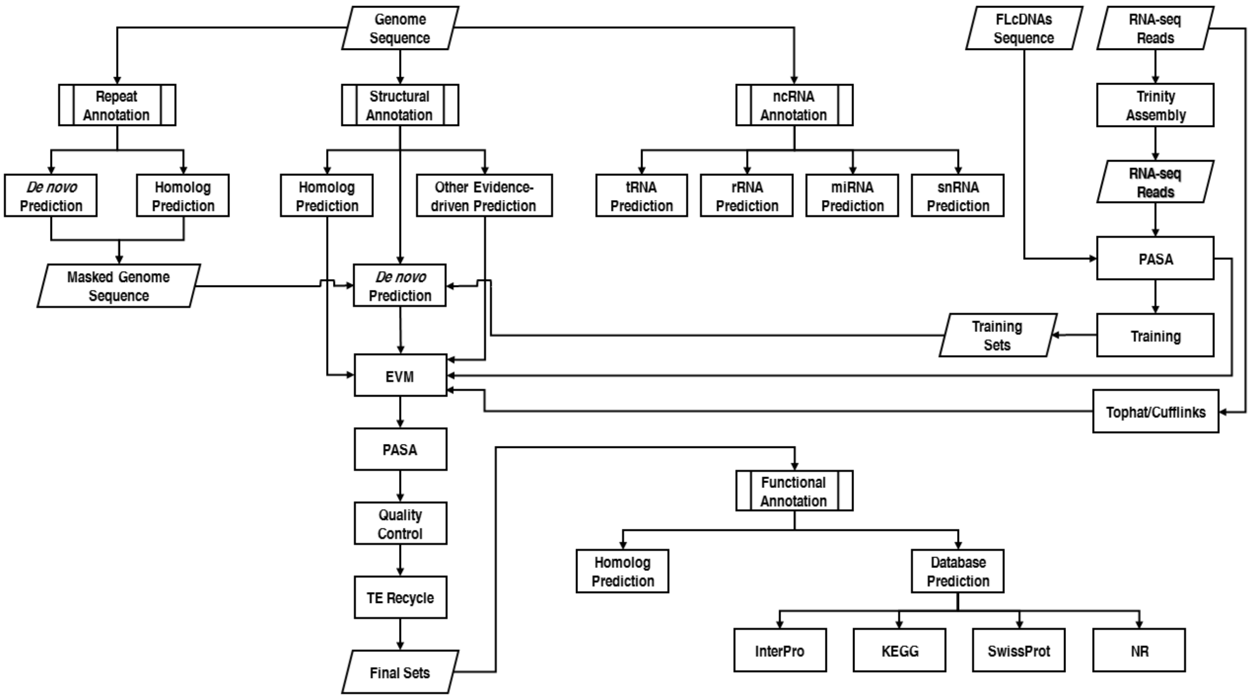


**Figure S1.** Protein-coding gene prediction process overview.

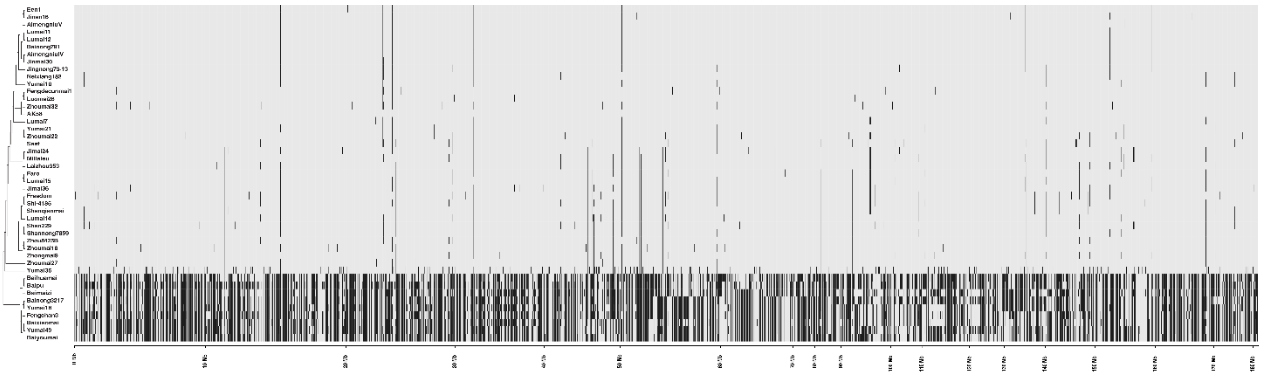

**Figure S2.** Neighbor-joining phylogenetic tree for SNP analysis of 36 1RS.1BL lines.

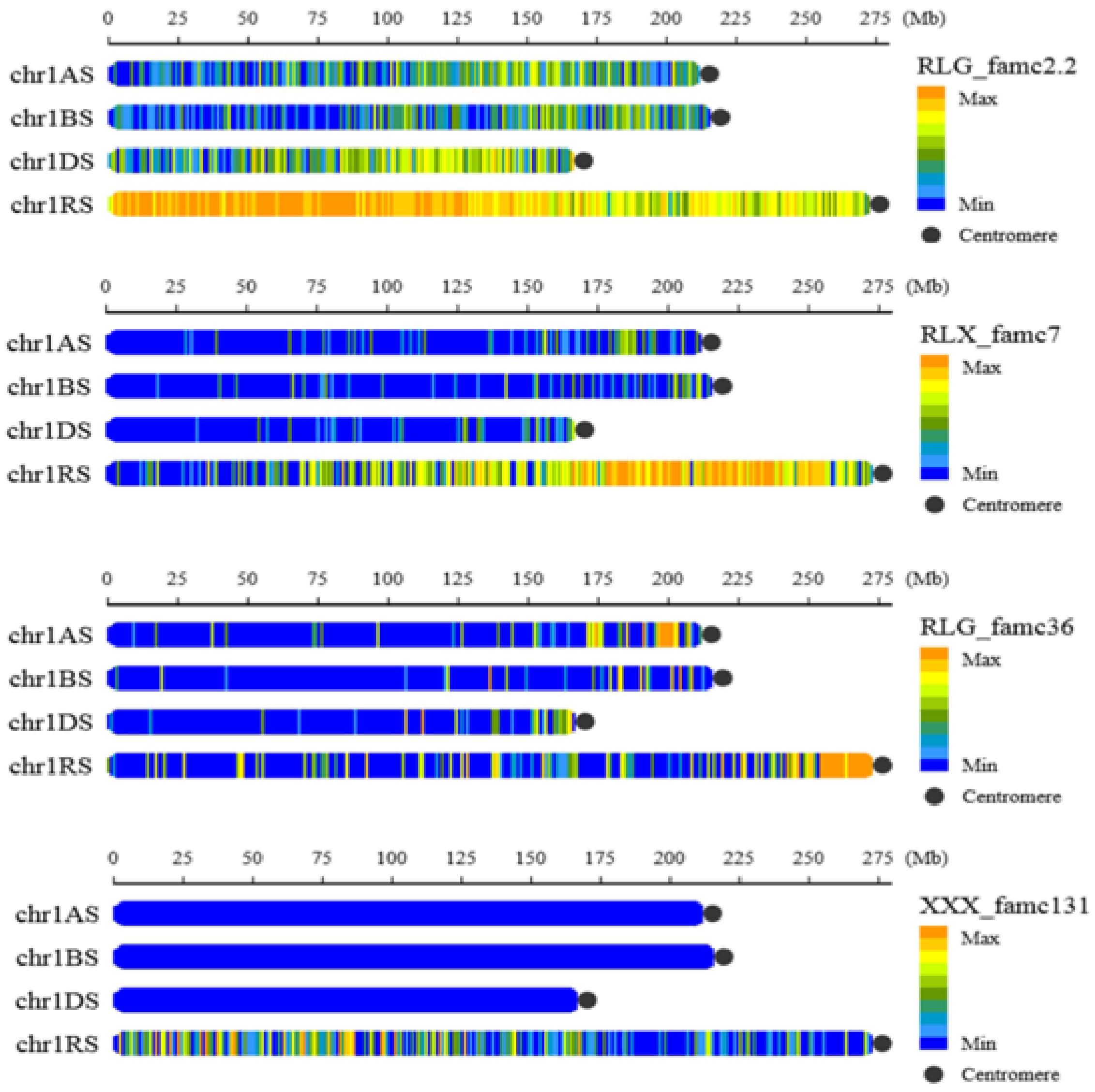


**Figure S3.** Distribution of some of the dominant retrotransposable elements in 1RS. The elements are identified on the right-hand side together with a color scale indicating the relative prominence of the elements.

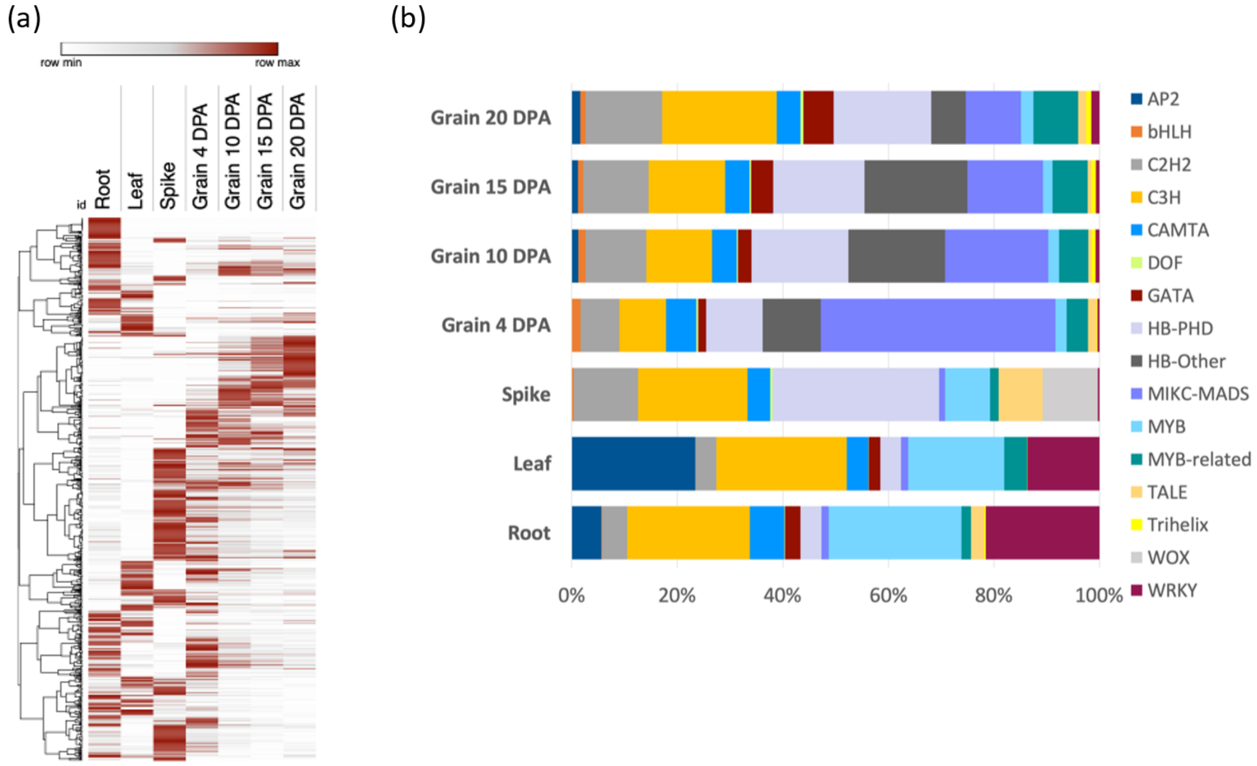


**Figure S4.** Transcript analysis. (a) Hierarchical clustering and relative gene expression patterns of 1RS gene models across the analyzed tissues. (b) Summary of levels of expression of the transcription factors. Decoding of the colours is indicated to the right of the panel.

**Table S1.** Pedigrees of wheat lines used for 660k SNP analysis.
**Table S2.** 1RS dominant TE information.
**Table S3.** PPR genes identified on the CS genome and their expression levels in different tissues.
**Table S4.** PPR genes located on the region of $Rf^{multi}$.
**Table S5.** PPR Genes identified on the AK58 genome and their expression levels in different tissues.
**Table S6.** Orthologous PPR genes from 1BS (CS) and 1RS (AK58).
**Table S7.** Orthologous genes from 1BS (CS) and 1RS (AK58) on the region of $Rf^{multi}/rf^{multi}$.
**Table S8.** PCR primers used in this study for subcellular localization.
**Table S9.** RGA Gene models analysis on the AK58 genome.
**Table S10.** A summary of the gene models in the morpheus clusters for contig-356, -445, -517 and -624 in the 1RS.1BL assembly.
