## Supplemental TableS1-9 for "1RS.1BL molecular resolution provides novel contributions to wheat improvement"

**Table S1 Pedigrees of wheat lines used for 660k SNP analysis.** The available pedigrees for the wheat lines used for the SNP analyses are summarized in the table. The different sources of rye genome introgressions are indicated in the document.

| Wheat variety | Pedigree | 1RS |
|---|---|---|
| Een1 | Lovrin10/761//Sumai3 | + |
| Jinan16 | Tal Shannongfu63/ Aimengniuv(Neuzucht) | + |
| Aimengniu V | Aifeng3//Mengxian201/Neuzucht | + |
| Lumai11 | Aimengniuiv(Neuzucht)/Shannongfu66 | + |
| Lumai12 | Lovrin10/Youbao//Ourou527/Aifeng3/3/Yexuan1 | + |
| Bainong791 | Bainong73_4262/Lovrin10//Bainong7122 | + |
| AimengniuIV | Aifeng3//Mengxian201/Neuzucht | + |
| Jinmai30 | Linfen5610/Shaan7587-5(Predgornia 2) | + |
| Jingnong79-13 | Youmanghong7/Lovrin10 | + |
| Neixiang182(Yumai17) | Yanda7406(rye and Kavkaz) /Nanyang75-6 | + |
| Yumai10 | Predgornia 2/Yanshi4 | + |
| Fengdecunmai1 | Zhou9811(hexaploid triticale)(/AK58(hexaploid triticale, Lovrin10) | + |
| Luomai26 | AK58(hexaploid triticale, Lovrin10)/Kaimai18 | + |
| Zhoumai32 | AK58(hexaploid triticale, Lovrin10)/Zhoumai24 | + |
| AK58 | Zhoumai11(hexaploid triticale)//Wenmai6(Lovrin10)/Zheng8960 | + |
| Lumai7 | Lovrin10/3/Virgilio/Ruluo//Youxuan26 | + |
| Yumai21 | Bainong791(Lovrin10)/Yumai2//Lumai1(Neuzucht)/Yanshi4 | + |
| Zhoumai22 | Zhoumai12(hexaploid triticale)/Yumai49(Lovrin10)//Zhoumai13(hexaploid triticale, Lovrin10) | + |
| Saet | – | + |

**Table S1 Pedigrees of wheat lines used for 660k SNP analysis.** The available pedigrees for the wheat lines used for the SNP analyses are summarized in the table. The different sources of rye genome introgressions are indicated in the document.

| **Wheat variety** | **Pedigree** | **1RS** | |
|---|---|---|---|
| Jimai24 | Anyang10/Aifeng1//75(89)(Lovrin10) | + | |
| Millaleu | – (Kavkaz) | + | |
| Laizhou953 | (Han5×Yexuan1)F3×7832H0—1 | + | |
| Faro | – | + | |
| Lumai15 | Tal/Yangmai1B1/Aimengniu Ⅱ (Neuzucht)//104-14 | + | |
| Jimai36 | Taishan5/shanqianmai(Predgornia 2)//Yannong78/3/Yuannong94354 | + | |
| Freedom | GR876(Kavkaz)/OH217=GR876/4/Logan*3/3/VA63-52-12/Logan/Blueboy | + | |
| Shi-4185 | Tal/Zhi8094(Predgornia 2)//Yumai2/3/Jimai26(Lovrin10) | + | |
| Shanqianmai(Predgornia 2) | Erythrospermum-315H60/Awnless 1 | + | |
| Lumai14 | C148(Lovrin13)/$F_4$-530 | + | |
| Shaan229 | Shaan7853( Kavkaz) //TB902/Xiaoyan6 | + | |
| Shaannong7859 | 7576$F_3$(Predgornia 2)/Shaan6861(2) | + | |
| Zhou8425B | Zhou78A(hexaploid triticale)/Annong7959 | + | |
| Zhongmai9 | Siyang936/83jian25(Predgornia 2) | + | |
| Zhoumai18 | Neixiang185(rye, Kavkaz and Lovrin10)/Zhoumai9(Lovrin10, Neuzucht) | + | |
| Zhoumai27 | Zhoumai16(hexaploid triticale, Lovrin10, Neuzucht)/AK58(hexaploid triticale, Lovrin10) | + | |
| Yumai35 | Mianyang84-27/Neixiang82C6(Lovrin10)// Yumai17(rye and Kavkaz) | + | |
| Baihuamai | – | – | Landrace |
| Baipu | – | – | Landrace |

**Table S1 Pedigrees of wheat lines used for 660k SNP analysis.** The available pedigrees for the wheat lines used for the SNP analyses are summarized in the table. The different sources of rye genome introgressions are indicated in the document.

| Wheat variety | Pedigree | 1RS | |
|---|---|---|---|
| Baimaizi | – | – | Landrace |
| Bainong3217 | Afu/Neixiang5//Xiannong39/3/3/Xinong64(4)3/Yanda24 | – | |
| Yumai18 | Zhengzhou761/Yanshi4 | – | |
| Fengchan3 | Danmai1/Xinong6028 | – | |
| Baixiaomai | – | – | Landrace |
| Yumai49 | Bainong791//Wenxuan1/Zhengzhou761/3/Yumai2 | – | |
| Baiyoumai | – | – | Landrace |

Reference: Zhuang QS. Chinese Wheat Improvement and Pedigree Analysis. Beijing: China Agriculture Press. 9(2003)

| | |
|---|---|
| Neuzucht | introduced from Germany (BRD), Petkus rye? |
| Lovrin10 | introduced from Romania, origin from Neuzucht*, Germany (BRD) |
| Lovrin13 | introduced from Romania, origin from Neuzucht*, Germany (BRD) |
| Kavkaz | introduced from Russia, origin from Neuzucht*, Germany (BRD) |
| Predgornia 2 | introduced from Russia, Predgornia 2 origin from Neuzucht, Germany (BRD) |

Reference: Rabinovich, SV. Importance of wheat-rye translocations for breeding modern cultivars of *Triticum aestivum L* . Euphytica. **100,** 323-340(1998).

| | |
|---|---|
| hexaploid triticale | origin from Guanghan Institute of Agricultural Sciences, Sichuan Province, China |
| rye | origin is not known |

**Table S2 1RS dominant TE information.**

| TE subfamily class | AK581AS | | | AK581RS | | | AK581DS | | | CS1BS | | |
|---|---|---|---|---|---|---|---|---|---|---|---|---|
| | Copy No. | Length | Ratio | Copy No. | Length | Ratio | Copy No. | Length | Ratio | Copy No. | Length | Ratio |
| LTR-Gypsy-RLG_famc2.2 | 1,225 | 1,531,692 | 1 | 11,942 | 13,628,604 | 5 | 1,371 | 2,149,538 | 1 | 1,050 | 1,176,055 | 1 |
| LTR-Gypsy-RLG_famc9.2 | 670 | 1,610,689 | 1 | 5,562 | 12,439,485 | 5 | 320 | 277,245 | 0 | 722 | 2,193,080 | 1 |
| LTR-Gypsy-RLG_famc2.5 | 697 | 525,232 | 0 | 3,045 | 3,311,348 | 1 | 685 | 503,762 | 0 | 738 | 497,822 | 0 |
| LTR-Gypsy-RLG_famc16 | 260 | 210,748 | 0 | 1,327 | 866,976 | 0 | 160 | 52,623 | 0 | 156 | 88,344 | 0 |
| LTR-Gypsy-RLG_famc36* | 202 | 238,016 | | 349 | 700,343 | 0 | 45 | 66,784 | 0 | 55 | 88,966 | 0 |
| LTR-Copia-RLC_famc12 | 198 | 80,478 | 0 | 527 | 234,214 | 0 | 144 | 69,029 | 0 | 187 | 78,056 | 0 |
| LTR-RLX_famc7 | 219 | 496,976 | | 3,828 | 8,703,915 | 3 | 147 | 263,487 | 0 | 188 | 394,317 | 0 |
| LTR-RLX_famc12 | 214 | 76,575 | 0 | 996 | 292,810 | 0 | 270 | 33,919 | 0 | 449 | 42,183 | 0 |
| LTR-RLX_famc21 | 28 | 26,024 | 0 | 307 | 200,346 | 0 | 1 | 989 | 0 | 72 | 54,965 | 0 |
| LINE-RIX_famc12 | 30 | 31,135 | 0 | 104 | 170,483 | 0 | 40 | 19,418 | 0 | 44 | 50,246 | 0 |
| Unknown-XXX_famc9 | 38 | 39,457 | 0 | 567 | 1,077,141 | 0 | 35 | 51,346 | 0 | 60 | 118,279 | 0 |
| Unknown-XXX_famc81 | 16 | 4,913 | 0 | 1,227 | 319,269 | 0 | 17 | 4,246 | 0 | 66 | 18,944 | 0 |
| Unknown-XXX_famc131 | 0 | 0 | 0 | 297 | 167,586 | 0 | 0 | 0 | 0 | 1 | 232 | 0 |
| Total | 3,797 | 4,871,935 | 2 | 30,078 | 42,112,520 | 15 | 3,235 | 3,492,386 | 2 | 3,788 | 4,801,489 | 2 |

**Table S2 1RS dominant TE information.**

| TE subfamily class | Ratio (length) | | | | Copy Number | | |
|---|---|---|---|---|---|---|---|
| | 1RS/1AS | 1RS/1BS | 1RS/1DS | Average | 1RS/1AS | 1RS/1BS | 1RS/1DS |
| LTR-Gypsy-RLG_famc2.2 | 9 | 12 | 6 | 9 | 9.7 | 11.4 | 8.7 |
| LTR-Gypsy-RLG_famc9.2 | 8 | 6 | 45 | 19 | 8.3 | 7.7 | 17.4 |
| LTR-Gypsy-RLG_famc2.5 | 6 | 7 | 7 | 7 | 4.4 | 4.1 | 4.4 |
| LTR-Gypsy-RLG_famc16 | 4 | 10 | 16 | 10 | 5.1 | 8.5 | 8.3 |
| LTR-Gypsy-RLG_famc36* | 3 | 8 | 10 | 7 | 1.7 | 6.3 | 7.8 |
| LTR-Copia-RLC_famc12 | 3 | 3 | 3 | 3 | 2.7 | 2.8 | 3.7 |
| LTR-RLX_famc7 | 18 | 22 | 33 | 24 | 17.5 | 20.4 | 26.0 |
| LTR-RLX_famc12 | 4 | 7 | 9 | 6 | 4.7 | 2.2 | 3.7 |
| LTR-RLX_famc21 | 8 | 4 | 203 | 71 | 11.0 | 4.3 | 307.0 |
| LINE-RIX_famc12 | 5 | 3 | 9 | 6 | 3.5 | 2.4 | 2.6 |
| Unknown-XXX_famc9 | 27 | 9 | 21 | 19 | 14.9 | 9.5 | 16.2 |
| Unknown-XXX_famc81 | 65 | 17 | 75 | 52 | 76.7 | 18.6 | 72.2 |
| Unknown-XXX_famc131 | - | - | - | - | - | - | - |
| Total | 160 | 108 | 437 | 233 | 160.1 | 98.1 | 478.0 |

Ratio: percentage of TE length/1AS or 1RS or 1DS or 1BS length

*RLG_Taes_Abia_consensus-1

**Table S3 PPR genes identified on the CS genome and their expression levels in different tissues.**

| Gene ID | Chr | Position(bp) | Class | PSI-mito[a] | Expression level(FPKM) | | | | | | |
|---|---|---|---|---|---|---|---|---|---|---|---|
| | | | | | Seedling leaf | Seedling root | FM | Grain 4 DPA | Grain 10 DPA | Grain 15 DPA | Grain 20 DPA |
| *TraesCS1B01G028800* | 1BS | 14,030,514 | PLS | 0.34594(0) | 0.10 | 0.33 | 0.64 | 0.59 | 0.36 | 0.32 | 0.31 |
| *TraesCS1B01G028900* | 1BS | 14,098,696 | PLS | 0.046995(1) | 0.04 | 0.04 | 0.33 | 0.26 | 0.14 | 0.15 | 0.10 |
| *TraesCS1B01G038200* | 1BS | 17,894,256 | 14-P | 0.66064(0) | 0.23 | 0.23 | 0.43 | 0.50 | 0.21 | 0.16 | 0.11 |
| *TraesCS1B01G038300* | 1BS | 17,964,601 | 10-P | 0.056598(1) | 0.07 | 0.07 | 0.64 | 0.16 | 0.08 | 0.01 | 0.01 |
| *TraesCS1B01G038400* | 1BS | 18,091,427 | 14-P | 0.015055(1) | 0.02 | 0.02 | 0.08 | 0.13 | 0.06 | 0.00 | 0.01 |
| *TraesCS1B01G038500* | 1BS | 18,116,277 | 15-P | 0.54383(0) | 0.04 | 0.04 | 0.40 | 0.35 | 0.16 | 0.10 | 0.05 |
| *TraesCS1B01G038600* | 1BS | 18,363,375 | 8-P | 0.014067(1) | 0.01 | 0.01 | 0.01 | 0.06 | 0.00 | 0.00 | 0.00 |
| *TraesCS1B01G039100* | 1BS | 18,683,381 | 16-P | 0.90407(0) | 0.00 | 0.00 | 0.08 | 0.14 | 0.05 | 0.02 | 0.01 |
| *TraesCS1B01G039200* | 1BS | 18,867,715 | 16-P | 0.59793(0) | 0.01 | 0.01 | 0.19 | 0.32 | 0.07 | 0.05 | 0.03 |
| *TraesCS1B01G039700* | 1BS | 19,073,838 | 15-P | 0.086936(0.9144) | 0.00 | 0.00 | 0.00 | 0.00 | 0.00 | 0.00 | 0.00 |
| *TraesCS1B01G052300* | 1BS | 32,592,265 | PLS | 0.69703(0) | 0.01 | 0.01 | 0.04 | 0.05 | 0.01 | 0.03 | 0.01 |
| *TraesCS1B01G066700* | 1BS | 50,987,332 | 3-P | 0.58922(0) | 0.18 | 0.18 | 4.83 | 1.86 | 1.00 | 0.45 | 0.67 |
| *TraesCS1B01G071600* | 1BS | 56,491,183 | 15-P | 0.474(0) | 0.12 | 0.12 | 0.55 | 0.59 | 0.23 | 0.10 | 0.08 |
| *TraesCS1B01G072300* | 1BS | 57,009,346 | 12-P | 0.71527(0) | 0.03 | 0.03 | 0.82 | 0.64 | 0.21 | 0.05 | 0.02 |
| *TraesCS1B01G072400* | 1BS | 57,016,242 | 14-P | 0.10571(0.0001) | 0.01 | 0.01 | 0.25 | 0.25 | 0.12 | 0.02 | 0.01 |
| *TraesCS1B01G072700* | 1BS | 57,104,191 | 11-P | 0.039141(1) | 0.00 | 0.00 | 0.12 | 0.10 | 0.02 | 0.00 | 0.02 |
| *TraesCS1B01G072900* | 1BS | 57,130,046 | 17-P | 0.86155(0) | 0.03 | 0.03 | 0.33 | 0.36 | 0.12 | 0.05 | 0.06 |
| *TraesCS1B01G073000* | 1BS | 57,138,687 | 4-P | 0.023775(1) | 0.00 | 0.00 | 0.00 | 0.00 | 0.00 | 0.00 | 0.00 |
| *TraesCS1B01G073100* | 1BS | 57,220,917 | 15-P | 0.009423(1) | 0.02 | 0.02 | 0.26 | 0.18 | 0.07 | 0.04 | 0.04 |

**Table S3 PPR genes identified on the CS genome and their expression levels in different tissues.**

| Gene ID | Chr | Position(bp) | Class | PSI-mito[a] | Expression level(FPKM) | | | | | | |
|---|---|---|---|---|---|---|---|---|---|---|---|
| | | | | | Seedling leaf | Seedling root | FM | Grain 4 DPA | Grain 10 DPA | Grain 15 DPA | Grain 20 DPA |
| *TraesCS1B01G073200* | 1BS | 57,374,298 | 5-P | 0.075852(1) | 0.21 | 0.21 | 0.34 | 1.39 | 0.71 | 0.26 | 0.21 |
| *TraesCS1B01G074600* | 1BS | 57,789,346 | 17-P | 0.92743(0) | 0.00 | 0.00 | 0.04 | 0.13 | 0.06 | 0.06 | 0.05 |
| *TraesCS1B01G075000* | 1BS | 58,114,326 | 15-P | 0.46132(0) | 0.01 | 0.01 | 0.16 | 0.19 | 0.15 | 0.01 | 0.00 |
| *TraesCS1B01G075300* | 1BS | 58,326,245 | 15-P | 0.10371(0.0008) | 0.00 | 0.00 | 0.08 | 0.05 | 0.05 | 0.00 | 0.04 |
| *TraesCS1B01G078800* | 1BS | 61,667,322 | 12-P | 0.011522(1) | 3.94 | 3.94 | 1.15 | 2.04 | 1.12 | 0.58 | 0.65 |
| *TraesCS1B01G084800* | 1BS | 69,547,272 | 11-P | 0.019015(1) | 0.00 | 0.00 | 0.00 | 0.00 | 0.00 | 0.00 | 0.00 |
| *TraesCS1B01G085300* | 1BS | 70,169,866 | 16-P | 0.41858(0) | 0.02 | 0.02 | 0.20 | 0.25 | 0.28 | 0.14 | 0.08 |
| *TraesCS1B01G085400* | 1BS | 70,185,997 | 16-P | 0.23027(1.89e-204) | 0.00 | 0.00 | 0.00 | 0.00 | 0.01 | 0.04 | 0.01 |
| *TraesCS1B01G085900* | 1BS | 70,333,799 | 15-P | 0.35558(0) | 0.03 | 0.03 | 0.20 | 0.28 | 0.12 | 0.07 | 0.08 |
| *TraesCS1B01G086000* | 1BS | 70,689,879 | 14-P | 0.17663(5.85e-97) | 0.00 | 0.00 | 0.04 | 0.00 | 0.01 | 0.00 | 0.00 |
| *TraesCS1B01G092500* | 1BS | 94,230,925 | 15-P | 0.63182(0) | 0.04 | 0.04 | 0.05 | 0.18 | 0.16 | 0.08 | 0.04 |
| *TraesCS1B01G092600* | 1BS | 94,325,745 | 10-P | 0.018009(1) | 0.12 | 0.12 | 0.48 | 0.32 | 0.42 | 0.52 | 0.49 |
| *TraesCS1B01G093500* | 1BS | 94,973,887 | 13-P | 0.6273(0) | 0.02 | 0.02 | 0.65 | 0.18 | 0.03 | 0.06 | 0.07 |
| *TraesCS1B01G108800* | 1BS | 120,884,346 | PLS | 0.6059(0) | 0.79 | 0.79 | 25.24 | 10.77 | 5.54 | 2.49 | 2.94 |
| *TraesCS1B01G109200* | 1BS | 121,013,048 | PLS | 0.27154(7.97e-287) | 0.11 | 0.11 | 0.43 | 0.31 | 0.15 | 0.01 | 0.02 |
| *TraesCS1B01G110500* | 1BS | 123,961,909 | PLS | 0.40904(0) | 0.04 | 0.04 | 0.49 | 0.21 | 0.12 | 0.04 | 0.04 |
| *TraesCS1B01G111700* | 1BS | 129,998,604 | 5-P | 0.10536(0) | 0.49 | 0.49 | 3.44 | 1.11 | 1.08 | 0.48 | 0.43 |
| *TraesCS1B01G111800* | 1BS | 130,001,406 | PLS | 0.034342(1) | 0.02 | 0.02 | 0.86 | 0.21 | 0.10 | 0.05 | 0.13 |
| *TraesCS1B01G122700* | 1BS | 148,410,844 | PLS | 0.085348(0.96) | 0.04 | 0.04 | 0.80 | 0.33 | 0.14 | 0.12 | 0.16 |

**Table S3 PPR genes identified on the CS genome and their expression levels in different tissues.**

| Gene ID | Chr | Position(bp) | Class | PSI-mito[a] | Expression level(FPKM) | | | | | | |
|---|---|---|---|---|---|---|---|---|---|---|---|
| | | | | | Seedling leaf | Seedling root | FM | Grain 4 DPA | Grain 10 DPA | Grain 15 DPA | Grain 20 DPA |
| *TraesCS1B01G136800* | 1BS | 174,762,914 | PLS | 0.027933(1) | 27.16 | 27.16 | 1.21 | 3.58 | 1.80 | 1.69 | 2.17 |
| *TraesCS1B01G136900* | 1BS | 174,783,888 | PLS | 0.031713(1) | 0.10 | 0.10 | 0.22 | 0.52 | 0.67 | 0.18 | 0.22 |
| *TraesCS1B01G137200* | 1BS | 175,275,522 | PLS | 0.11592(7.03e-11) | 5.13 | 5.13 | 13.90 | 3.52 | 1.83 | 1.67 | 1.76 |
| *TraesCS1B01G137400* | 1BS | 175,428,009 | 9-P | 0.017238(1) | 0.16 | 0.16 | 2.25 | 0.91 | 0.82 | 0.78 | 0.57 |
| *TraesCS1B01G140500* | 1BS | 186,518,365 | 3-P | 0.010863(1) | 0.01 | 0.01 | 0.49 | 0.59 | 0.14 | 0.27 | 0.05 |
| *TraesCS1B01G150100* | 1BS | 229,237,139 | 7-P | 0.0066905(1) | 15.70 | 15.70 | 1.86 | 4.30 | 1.20 | 0.89 | 0.50 |

[a]estimation of mitochondria location signal value

**Table S4 PPR genes located on the region of *Rf*$^{multi}$.**

| Gene/Marker_ID | Chr | Length | Start | End | Direct |
|---|---|---|---|---|---|
| *TraesCS1B01G028800* | 1BS | 1122 | 14,030,514 | 14,031,722 | - |
| *TraesCS1B01G028900* | 1BS | 1692 | 14,098,696 | 14,100,474 | - |
| *Xucr_8* | 1BS | 1437 | 15,288,153 | 15,289,589 | + |
| *TraesCS1B01G038200* | 1BS | 2309 | 17,894,256 | 17,896,564 | - |
| *TraesCS1B01G038300* | 1BS | 1641 | 17,964,601 | 17,966,241 | - |
| *TraesCS1B01G038400* | 1BS | 1827 | | 18,093,253 | - |
| *TraesCS1B01G038500* | 1BS | 2373 | 18,116,277 | 18,118,649 | - |
| *TraesCS1B01G038600* | 1BS | 1092 | | 18,364,466 | - |
| *TraesCS1B01G039100* | 1BS | 2352 | 18,683,381 | 18,685,732 | + |
| *TraesCS1B01G039200* | 1BS | 2355 | 18,867,715 | 18,870,069 | + |
| *TraesCS1B01G039700* | 1BS | 1827 | 19,073,838 | 19,075,664 | - |
| *TraesCS1B01G052300* | 1BS | 1458 | 32,592,265 | 32,594,796 | - |
| *TraesCS1B01G066700* | 1BS | 810 | 50,987,332 | 50,988,141 | + |
| *TraesCS1B01G071600* | 1BS | 2647 | 56,491,183 | 56,493,829 | - |
| *TraesCS1B01G072300* | 1BS | 1782 | 57,009,346 | 57,011,127 | + |
| *TraesCS1B01G072400* | 1BS | 2088 | 57,016,242 | 57,018,329 | + |
| *TraesCS1B01G072700* | 1BS | 1533 | 57,104,191 | 57,105,723 | + |
| *TraesCS1B01G072900* | 1BS | 2622 | 57,130,046 | 57,132,667 | + |
| *TraesCS1B01G073000* | 1BS | 786 | 57,138,687 | 57,139,472 | + |
| *TraesCS1B01G073100* | 1BS | 1821 | 57,220,917 | 57,222,737 | + |

**Table S4 PPR genes located on the region of *Rf*$^{multi}$.**

| Gene/Marker_ID | Chr | Length | Start | End | Direct |
|---|---|---|---|---|---|
| *TraesCS1B01G073200* | 1BS | 912 | 57,374,298 | 57,375,209 | + |
| *TraesCS1B01G074600* | 1BS | 2343 | 57,789,346 | 57,791,688 | - |
| *TraesCS1B01G075000* | 1BS | 1995 | 58,114,326 | 58,116,320 | - |
| *TraesCS1B01G075300* | 1BS | 1794 | 58,326,245 | 58,328,038 | - |
| *Xucr_5* | 1BS | 761 | 59,833,238 | 59,833,998 | - |
| *TraesCS1B01G078800* | 1BS | 1662 | 61,667,322 | 61,668,983 | + |
| *TraesCS1B01G084800* | 1BS | 1395 | 69,547,272 | 69,548,666 | - |
| *TraesCS1B01G085300* | 1BS | 2379 | 70,169,866 | 70,172,244 | + |
| *TraesCS1B01G085400* | 1BS | 1926 | 70,185,997 | 70,187,922 | + |
| *TraesCS1B01G085900* | 1BS | 2445 | 70,333,799 | 70,336,243 | + |
| *TraesCS1B01G086000* | 1BS | 2061 | 70,689,879 | 70,696,184 | + |
| *Xucr_3* | 1BS | 588 | 84,962,211 | 84,962,799 | + |
| *Xucr_4* | 1BS | 500 | 91,556,341 | 91,557,658 | + |
| *TraesCS1B01G092500* | 1BS | 2352 | 94,230,925 | 94,233,276 | - |
| *TraesCS1B01G092600* | 1BS | 1554 | 94,325,745 | 94,327,298 | - |
| *TraesCS1B01G093500* | 1BS | 1914 | 94,973,887 | 94,975,800 | + |
| *TraesCS1B01G108800* | 1BS | 1359 | 120,884,346 | 120,888,721 | - |
| *TraesCS1B01G109200* | 1BS | 1812 | 121,013,048 | 121,014,970 | - |
| *TraesCS1B01G110500* | 1BS | 1863 | 123,961,909 | 123,963,771 | + |
| *TraesCS1B01G111700* | 1BS | 1083 | 129,998,604 | 129,999,686 | + |

**Table S4 PPR genes located on the region of *Rf*$^{multi}$.**

| Gene/Marker_ID | Chr | Length | Start | End | Direct |
|---|---|---|---|---|---|
| *TraesCS1B01G111800* | 1BS | 1080 | 130,001,406 | 130,002,485 | + |
| *TraesCS1B01G122700* | 1BS | 1473 | 148,410,844 | 148,412,316 | + |
| *TraesCS1B01G136800* | 1BS | 649 | 174,762,914 | 174,763,562 | - |
| *TraesCS1B01G136900* | 1BS | 378 | 174,783,888 | 174,784,265 | - |
| *TraesCS1B01G137200* | 1BS | 336 | 175,275,522 | 175,275,857 | - |
| *TraesCS1B01G137400* | 1BS | 1647 | 175,428,009 | 175,429,655 | - |
| *TraesCS1B01G140500* | 1BS | 798 | 186,518,365 | 186,519,267 | + |
| *TraesCS1B01G150100* | 1BS | 3453 | 229,237,139 | 229,275,531 | + |

**Table S5 PPR genes identified on the AK58 genome and their expression levels in different tissues.**

| Gene ID | Chr | Position(bp) | Class | PSI-mito[a] | Expression level(FPKM) | | | | | | |
|---|---|---|---|---|---|---|---|---|---|---|---|
| | | | | | Seedling leaf | Seedling root | FM | Grain 4 DPA | Grain 10 DPA | Grain 15 DPA | Grain 20 DPA |
| *Xucr_8* | 1RS | 585,950 | | | | | | | | | |
| *TraesAK58CH1B01G001600* | 1RS | 1,362,618 | PLS | 0.15938(4.41e-66) | 0.00 | 0.04 | 0.01 | 0.09 | 0.09 | 0.16 | 0.01 |
| *TraesAK58CH1B01G041500* | 1RS | 44,460,287 | 2-P | 0.83096(0) | 0.06 | 0.51 | 2.27 | 1.10 | 0.62 | 0.30 | 0.37 |
| *TraesAK58CH1B01G043600* | 1RS | 48,608,965 | 14-P | 0.014923(1) | 0.10 | 0.13 | 0.02 | 0.24 | 0.22 | 0.09 | 0.03 |
| *TraesAK58CH1B01G043700* | 1RS | 48,622,083 | 14-P | 0.76317(0) | 0.00 | 0.00 | 0.00 | 0.00 | 0.00 | 0.00 | 0.00 |
| *TraesAK58CH1B01G043800* | 1RS | 48,672,479 | 14-P | 0.11632(3.5e-11) | 0.17 | 0.23 | 0.15 | 0.48 | 0.46 | 0.31 | 0.25 |
| *TraesAK58CH1B01G043900* | 1RS | 48,697,960 | 15-P | 0.76866(0) | 0.01 | 0.04 | 0.29 | 0.36 | 0.11 | 0.13 | 0.07 |
| *TraesAK58CH1B01G045100* | 1RS | 49,148,431 | 17-P | 0.84539(0) | 0.00 | 0.00 | 0.07 | 0.10 | 0.02 | 0.05 | 0.01 |
| *TraesAK58CH1B01G045250* | 1RS | 49,407,657 | 17-P | 0.097262(0.078) | 0.00 | 0.00 | 0.00 | 0.00 | 0.00 | 0.00 | 0.00 |
| *Xucr_5* | 1RS | 52,342,853 | | | | | | | | | |
| *TraesAK58CH1B01G054300* | 1RS | 62,878,940 | 17-P | 0.76681(0) | 0.00 | 0.02 | 0.12 | 0.24 | 0.08 | 0.06 | 0.02 |
| *TraesAK58CH1B01G054600* | 1RS | 63,198,799 | 15-P | 0.82319(0) | 0.00 | 0.10 | 0.13 | 0.19 | 0.10 | 0.12 | 0.06 |
| *TraesAK58CH1B01G054700* | 1RS | 63,286,316 | 16-P | 0.79683(0) | 0.00 | 0.03 | 0.10 | 0.03 | 0.05 | 0.00 | 0.06 |
| *TraesAK58CH1B01G054800* | 1RS | 63,329,752 | 15-P | 0.79944(0) | 0.00 | 0.01 | 0.06 | 0.03 | 0.01 | 0.00 | 0.03 |
| *TraesAK58CH1B01G062800* | 1RS | 81,045,350 | 13-P | 0.76962(0) | 0.03 | 0.18 | 0.18 | 0.12 | 0.05 | 0.04 | 0.07 |
| *TraesAK58CH1B01G075300* | 1RS | 104,250,542 | PLS | 0.77825(0) | 0.98 | 7.25 | 25.08 | 14.39 | 12.75 | 8.22 | 5.99 |
| *TraesAK58CH1B01G075700* | 1RS | 104,650,641 | PLS | 0.53681(0) | 0.10 | 0.59 | 0.44 | 0.94 | 0.47 | 0.39 | 0.39 |
| *TraesAK58CH1B01G077300* | 1RS | 109,429,796 | PLS | 0.47379(0) | 0.03 | 0.12 | 0.32 | 0.29 | 0.22 | 0.23 | 0.16 |
| *TraesAK58CH1B01G077800* | 1RS | 110,382,945 | 4-P | 0.37101(0) | 0.31 | 0.31 | 2.91 | 0.80 | 0.67 | 0.67 | 0.63 |

**Table S5 PPR genes identified on the AK58 genome and their expression levels in different tissues.**

| Gene ID | Chr | Position(bp) | Class | PSI-mito[a] | Expression level(FPKM) | | | | | | |
|---|---|---|---|---|---|---|---|---|---|---|---|
| | | | | | Seedling leaf | Seedling root | FM | Grain 4 DPA | Grain 10 DPA | Grain 15 DPA | Grain 20 DPA |
| *TraesAK58CH1B01G077900* | 1RS | 110,399,492 | PLS | 0.15402(3.0e-57) | 0.01 | 0.14 | 0.52 | 0.34 | 0.25 | 0.14 | 0.30 |
| *TraesAK58CH1B01G084800* | 1RS | 123,613,223 | PLS | 0.051203(1) | 0.01 | 0.06 | 0.35 | 0.18 | 0.18 | 0.06 | 0.10 |
| *TraesAK58CH1B01G097400* | 1RS | 149,960,198 | PLS | 0.036487(1) | 15.31 | 21.74 | 2.31 | 0.32 | 0.08 | 0.44 | 0.23 |
| *TraesAK58CH1B01G097500* | 1RS | 150,239,425 | PLS | 0.026069(1) | 2.11 | 2.46 | 1.35 | 1.56 | 0.83 | 0.58 | 0.45 |
| *TraesAK58CH1B01G097700* | 1RS | 150,786,022 | PLS | 0.025267(1) | 1.04 | 15.58 | 3.01 | 1.81 | 1.63 | 2.83 | 1.74 |
| *TraesAK58CH1B01G097900* | 1RS | 151,065,049 | 9-P | 0.036791(1) | 0.16 | 0.46 | 1.03 | 0.39 | 0.60 | 0.45 | 0.53 |
| *TraesAK58CH1B01G104100* | 1RS | 162,021,582 | 14-P | 0.27077(2.4e-285) | 0.49 | 0.89 | 1.43 | 2.57 | 1.70 | 1.40 | 1.00 |

[a]estimation of mitochondria location signal value

**Table S6 Orthologous PPR genes from 1BS (CS) and 1RS (AK58).**

| AK58_PPR | | | | | CS_PPR | | | | | |
|---|---|---|---|---|---|---|---|---|---|---|
| Gene ID | Chr | Position | PSI-mito[a] | P motif number | Gene ID | Chr | Position | PSI-mito[a] | P motif number | Class |
| *TraesAK58CH1B01G001600* | 1RS | 1,362,618 | 0.15938(4.4e-66) | PLS | *TraesCS1B01G028900* | 1BS | 14,098,696 | 0.046995(1) | PLS | PLS |
| *TraesAK58CH1B01G041500* | 1RS | 44,460,287 | 0.83096(0) | 2-P | *TraesCS1B01G066700* | 1BS | 50,987,332 | 0.58922(0) | 3-P | P |
| *TraesAK58CH1B01G043800* | 1RS | 48,672,479 | 0.11632(3.5e-11) | 14-P | *TraesCS1B01G071600* | 1BS | 56,491,183 | 0.474(0) | 15-P | P |
| *TraesAK58CH1B01G043900* | 1RS | 48,697,960 | 0.76866(0) | 15-P | *TraesCS1B01G074600* | 1BS | 57,791,688 | 0.92743(0) | 17-P | P |
| *TraesAK58CH1B01G045100* | 1RS | 49,148,431 | 0.84539(0) | 17-P | *TraesCS1B01G072300* | 1BS | 57,009,346 | 0.71527(0) | 12-P | P |
| *TraesAK58CH1B01G045250* | 1RS | 49,407,657 | 0.097262(0.078) | 17-P | *TraesCS1B01G072900* | 1BS | 57,130,046 | 0.86155(0) | 17-P | P |
| *TraesAK58CH1B01G045250* | 1RS | 49,407,657 | 0.097262(0.078) | 17-P | *TraesCS1B01G074600* | 1BS | 57,789,346 | 0.92743(0) | 17-P | P |
| *TraesAK58CH1B01G054300* | 1RS | 62,878,940 | 0.76681(0) | 17-P | *TraesCS1B01G085300* | 1BS | 70,169,866 | 0.41858(0) | 16-P | P |
| *TraesAK58CH1B01G054600* | 1RS | 63,198,799 | 0.82319(0) | 15-P | *TraesCS1B01G085900* | 1BS | 70,333,799 | 0.35558(0) | 15-P | P |
| *TraesAK58CH1B01G062800* | 1RS | 81,045,350 | 0.76962(0) | 13-P | *TraesCS1B01G093500* | 1BS | 94,973,887 | 0.6273(0) | 13-P | P |
| *TraesAK58CH1B01G075300* | 1RS | 104,250,542 | 0.77825(0) | PLS | *TraesCS1B01G108800* | 1BS | 120,884,346 | 0.6059(0) | PLS | PLS |
| *TraesAK58CH1B01G075700* | 1RS | 104,650,641 | 0.53681(0) | PLS | *TraesCS1B01G109200* | 1BS | 121,013,048 | 0.27154(8.0e-287) | PLS | PLS |
| *TraesAK58CH1B01G077300* | 1RS | 109,429,796 | 0.47379(0) | PLS | *TraesCS1B01G110500* | 1BS | 123,961,909 | 0.40904(0) | PLS | PLS |
| *TraesAK58CH1B01G077800* | 1RS | 110,382,945 | 0.37101(0) | 4-P | *TraesCS1B01G111700* | 1BS | 129,998,604 | 0.10536(0) | 5-P | P |
| *TraesAK58CH1B01G077900* | 1RS | 110,399,492 | 0.15402(2.9e-57) | PLS | *TraesCS1B01G111800* | 1BS | 130,001,406 | 0.034342(1) | PLS | PLS |
| *TraesAK58CH1B01G084800* | 1RS | 123,613,223 | 0.051203(1) | PLS | *TraesCS1B01G122700* | 1BS | 148,410,844 | 0.085348(0.96) | PLS | PLS |
| *TraesAK58CH1B01G097400* | 1RS | 149,960,198 | 0.036487(1) | PLS | *TraesCS1B01G136800* | 1BS | 174,762,913 | 0.027933(1) | PLS | PLS |
| *TraesAK58CH1B01G097500* | 1RS | 150,239,425 | 0.026069(1) | PLS | *TraesCS1B01G136900* | 1BS | 174,783,888 | 0.031713(1) | PLS | PLS |
| *TraesAK58CH1B01G097700* | 1RS | 150,786,022 | 0.025267(1) | PLS | *TraesCS1B01G137200* | 1BS | 175,275,522 | 0.11592(7.03e-11) | PLS | PLS |

**Table S6 Orthologous PPR genes from 1BS (CS) and 1RS (AK58).**

| AK58_PPR | | | | | CS_PPR | | | | | |
|---|---|---|---|---|---|---|---|---|---|---|
| **Gene ID** | **Chr** | **Position** | **PSI-mito[a]** | **P motif number** | **Gene ID** | **Chr** | **Position** | **PSI-mito[a]** | **P motif number** | **Class** |
| *TraesAK58CH1B01G097900* | 1RS | 151,065,049 | 0.036791(1) | 9-P | *TraesCS1B01G137400* | 1BS | 175,428,009 | 0.017238(1) | 9-P | P |
| *TraesAK58CH1B01G104100* | 1RS | 162,021,582 | 0.27077(2.4e-285) | 14-P | *TraesCS1B01G140500* | 1BS | 186,518,365 | 0.010863(1) | 3-P | P |

[a]estimation of mitochondria location signal value

**Table S7 Orthologous genes from 1BS (CS) and 1RS (AK58) on the region of *Rf*$^{multi}$.**

| CS_PPR | | | | | AK58_PPR | | | | |
|---|---|---|---|---|---|---|---|---|---|
| **start** | **end** | **Chr** | **Direct** | **Gene_ID** | **Gene_ID** | **Chr** | **start** | **end** | **Direct** |
| 56,310,657 | 56,314,832 | 1BS | + | *TraesCS1B01G071400* | *TraesAK58CH1B01G043400* | 1RS | 48,461,156 | 48,466,866 | + |
| 56,316,840 | 56,322,275 | 1BS | - | *TraesCS1B01G071500* | *TraesAK58CH1B01G043500* | 1RS | 48,467,609 | 48,474,219 | - |
| 56,491,183 | 56,493,829 | 1BS | - | *TraesCS1B01G071600* | | | | | |
| 56,994,218 | 57,003,342 | 1BS | + | *TraesCS1B01G072200* | *TraesAK58CH1B01G043600* | 1RS | 48,608,965 | 48,610,788 | - |
| 56,976,659 | 56,977,645 | 1BS | | *TraesCS1B01G072100* | *TraesAK58CH1B01G043700* | 1RS | 48,622,083 | 48,628,102 | - |
| 56,890,943 | 56,894,269 | 1BS | - | *TraesCS1B01G072000* | *TraesAK58CH1B01G043800* | 1RS | 48,672,479 | 48,675,076 | - |
| 56,883,391 | 56,884,076 | 1BS | | *TraesCS1B01G071900* | ***TraesAK58CH1B01G043900*** | 1RS | 48,697,960 | 48,700,407 | - |
| 56,883,314 | 56,888,127 | 1BS | + | *TraesCS1B01G071800* | | | | | |
| 56,879,177 | 56,883,068 | 1BS | - | *TraesCS1B01G071700* | *TraesAK58CH1B01G044000* | 1RS | 48,804,182 | 48,811,468 | - |
| 57,009,346 | 57,011,127 | 1BS | + | ***TraesCS1B01G072300*** | *TraesAK58CH1B01G044100* | 1RS | 48,813,836 | 48,815,083 | - |
| 57,016,242 | 57,018,329 | 1BS | + | *TraesCS1B01G072400* | *TraesAK58CH1B01G044200* | 1RS | 48,819,935 | 48,820,870 | + |
| 57,022,598 | 57,024,451 | 1BS | - | *TraesCS1B01G072500* | | | | | |
| 57,044,480 | 57,051,132 | 1BS | - | *TraesCS1B01G072600* | | | | | |
| 57,104,191 | 57,105,723 | 1BS | + | *TraesCS1B01G072700* | *TraesAK58CH1B01G044300* | 1RS | 48,849,495 | 48,850,496 | + |
| 57,110,355 | 57,110,917 | 1BS | - | *TraesCS1B01G072800* | | | | | |
| 57,130,046 | 57,132,667 | 1BS | + | ***TraesCS1B01G072900*** | *TraesAK58CH1B01G044400* | 1RS | 48,864,756 | 48,865,775 | + |
| 57,138,687 | 57,139,472 | 1BS | + | *TraesCS1B01G073000* | | | | | |
| 57,220,917 | 57,222,737 | 1BS | + | *TraesCS1B01G073100* | *TraesAK58CH1B01G044500* | 1RS | 48,909,951 | 48,910,844 | + |
| 57,374,298 | 57,375,209 | 1BS | + | *TraesCS1B01G073200* | *TraesAK58CH1B01G044600* | 1RS | 48,932,924 | 48,933,925 | + |

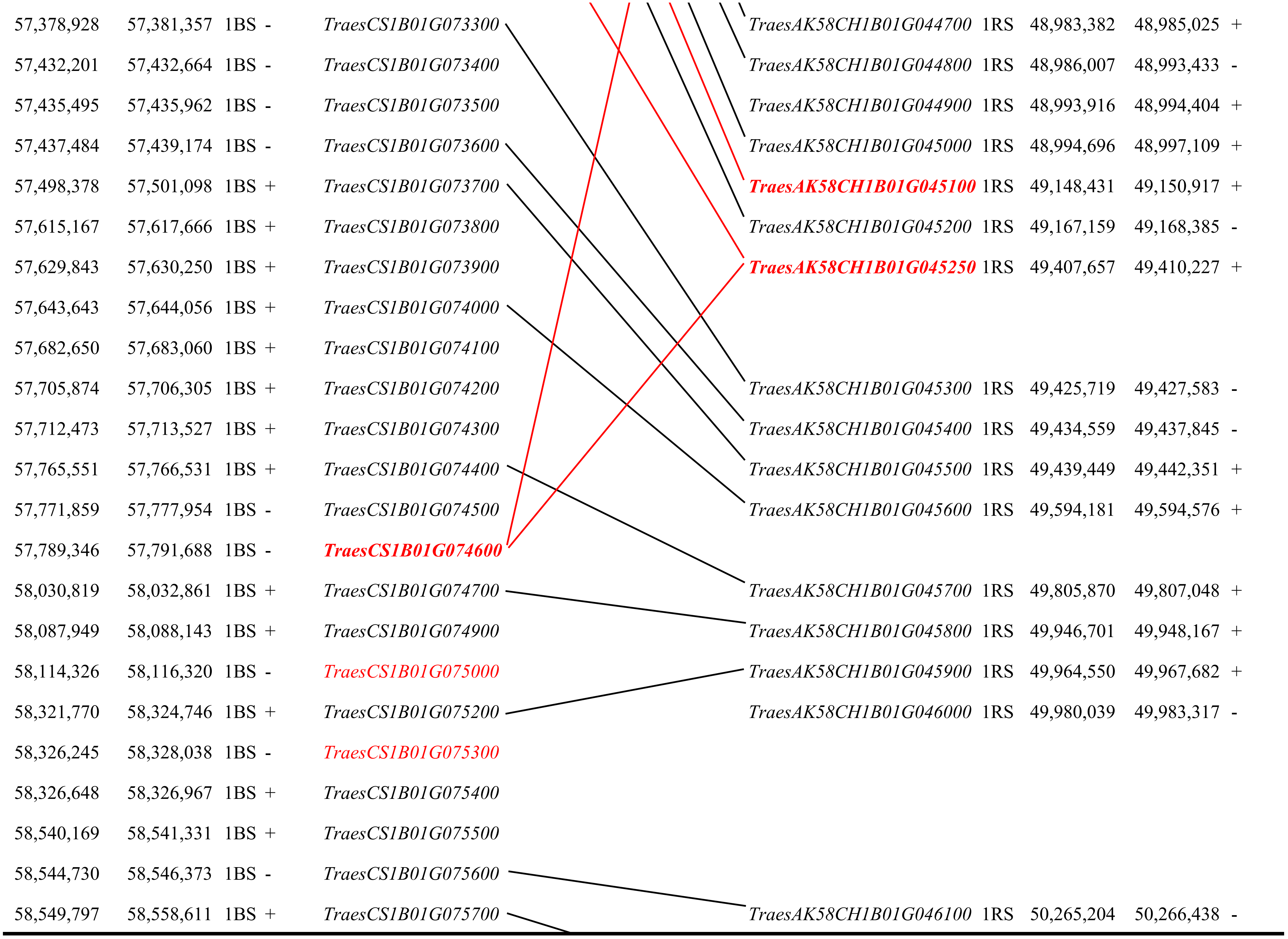
57,378,928 57,381,357 1BS - TraesCS1B01G073300
57,432,201 57,432,664 1BS - TraesCS1B01G073400
57,435,495 57,435,962 1BS - TraesCS1B01G073500
57,437,484 57,439,174 1BS - TraesCS1B01G073600
57,498,378 57,501,098 1BS + TraesCS1B01G073700
57,615,167 57,617,666 1BS + TraesCS1B01G073800
57,629,843 57,630,250 1BS + TraesCS1B01G073900
57,643,643 57,644,056 1BS + TraesCS1B01G074000
57,682,650 57,683,060 1BS + TraesCS1B01G074100
57,705,874 57,706,305 1BS + TraesCS1B01G074200
57,712,473 57,713,527 1BS + TraesCS1B01G074300
57,765,551 57,766,531 1BS + TraesCS1B01G074400
57,771,859 57,777,954 1BS - TraesCS1B01G074500
57,789,346 57,791,688 1BS - TraesCS1B01G074600
58,030,819 58,032,861 1BS + TraesCS1B01G074700
58,087,949 58,088,143 1BS + TraesCS1B01G074900
58,114,326 58,116,320 1BS - TraesCS1B01G075000
58,321,770 58,324,746 1BS + TraesCS1B01G075200
58,326,245 58,328,038 1BS - TraesCS1B01G075300
58,326,648 58,326,967 1BS + TraesCS1B01G075400
58,540,169 58,541,331 1BS + TraesCS1B01G075500
58,544,730 58,546,373 1BS - TraesCS1B01G075600
58,549,797 58,558,611 1BS + TraesCS1B01G075700
TraesAK58CH1B01G044700 1RS 48,983,382 48,985,025 +
TraesAK58CH1B01G044800 1RS 48,986,007 48,993,433 -
TraesAK58CH1B01G044900 1RS 48,993,916 48,994,404 +
TraesAK58CH1B01G045000 1RS 48,994,696 48,997,109 +
TraesAK58CH1B01G045100 1RS 49,148,431 49,150,917 +
TraesAK58CH1B01G045200 1RS 49,167,159 49,168,385 -
TraesAK58CH1B01G045250 1RS 49,407,657 49,410,227 +
TraesAK58CH1B01G045300 1RS 49,425,719 49,427,583 -
TraesAK58CH1B01G045400 1RS 49,434,559 49,437,845 -
TraesAK58CH1B01G045500 1RS 49,439,449 49,442,351 +
TraesAK58CH1B01G045600 1RS 49,594,181 49,594,576 +
TraesAK58CH1B01G045700 1RS 49,805,870 49,807,048 +
TraesAK58CH1B01G045800 1RS 49,946,701 49,948,167 +
TraesAK58CH1B01G045900 1RS 49,964,550 49,967,682 +
TraesAK58CH1B01G046000 1RS 49,980,039 49,983,317 -
TraesAK58CH1B01G046100 1RS 50,265,204 50,266,438 -

| | | | | |
|---|---|---|---|---|
| 58,563,500 | 58,586,329 | 1BS | + | *TraesCS1B01G075800* |
| 58,705,113 | 58,709,166 | 1BS | + | *TraesCS1B01G075900* |
| 58,752,977 | 58,754,134 | 1BS | + | *TraesCS1B01G076000* |

| | | | | |
|---|---|---|---|---|
| *TraesAK58CH1B01G046200* | 1RS | 50,268,538 | 50,270,472 | - |
| *TraesAK58CH1B01G046300* | 1RS | 50,345,169 | 50,345,543 | + |
| *TraesAK58CH1B01G046400* | 1RS | 50,348,134 | 50,359,423 | + |

Notes: red indicate PPR genes

**Table S8 PCR primers used in this study for subcellular localization.**

| Primer ID | Primer sequence (5' to 3') | Application |
|---|---|---|
| *TraesAK58CH1B01G045100* cDNAF | GCTTGATTGAGTGACGGGAAGA | *TraesAK58CH1B01G045100* cDNA cloning from total RNA |
| *TraesAK58CH1B01G045100* cDNAR | GCGTCGCTAGGTATTTCG | |
| *TraesCS1B01G072900* cDNAF | GGCTGTGCAACTCCAAACTCT | *TraesCS1B01G072900* cDNA cloning from total RNA |
| *TraesCS1B01G072900* cDNAR | CAATTATACTGGCAGGAGGGCT | |
| *TraesCS1B01G072300* cDNAF | TCCTCCTCCTAGCCGATG | *TraesCS1B01G072300* cDNA cloning from total RNA |
| *TraesCS1B01G072300* cDNAR | AGCTGGGGACTATACCAGTGA | |
| *TraesCS1B01G074600* cDNAF | CCTCTGCCTCCGCACTACTC | *TraesCS1B01G074600* cDNA cloning from total RNA |
| *TraesCS1B01G074600* cDNAR | GAAGTAGTTGAAGCTTCAAGTAGCA | |
| *TraesAK58CH1B01G045100* F | GGGACTCTAGAggatccATGTCCCGCCTCCGCCGC | GFP-fusion for subcellular localization |
| *TraesAK58CH1B01G045100* R | CACCATGGTACCgtcgacGCCAAATTCACCAAAA | |
| *TraesCS1B01G072900* F | GGGACTCTAGAggatccATGTCCCGCCGCATCCGC | |
| *TraesCS1B01G072900* R | CACCATGGTACCgtcgacGACAACTAATCCGTCGAA | |
| *TraesCS1B01G072300* F | GGGACTCTAGAggatccATGTCGGGCCTCCGTCTC | |
| *TraesCS1B01G072300* R | CACCATGGTACCgtcgacTACCAGTGATTCCATGGCATT | |
| *TraesCS1B01G074600* F | GGGACTCTAGAggatccATGTCCCGCCTCCGCC | |
| *TraesCS1B01G074600* R | CACCATGGTACCgtcgacACAAATAAAGTCCGGCCAT | |

**Table S9 RGA gene models analysis on the AK58 genome.**

| Chr | Unique to AK58 | Start | Domain | Class | absent from 1BS CS | domain from Phyre2 analysis, 3D fold for domain |
|---|---|---|---|---|---|---|
| 1B | *TraesAK58CH1B01G001310* | 1,281,152 | LRR,NB-ARC | NL | | no significant hit |
| 1B | *TraesAK58CH1B01G004400* | 2,589,629 | CC,LRR,NB-ARC | CNL | | disease resistance rpp13-like protein 4, c6j5tC_ |
| 1B | *TraesAK58CH1B01G005100* | 2,835,356 | STTK,TM | RLK | | tyrosine-protein kinase abl1, c2fo0A_ |
| 1B | *TraesAK58CH1B01G005800* | 3,096,772 | STTK,TM | RLK | | tyrosine-protein kinase abl1, c2fo0A_ |
| 1B | *TraesAK58CH1B01G005900* | 3,154,442 | CC,LRR,NB-ARC | CNL | | disease resistance rpp13-like protein 4, c6j5tC_ |
| 1B | *TraesAK58CH1B01G006500* | 3,784,310 | | RLK | | tyrosine-protein kinase abl1, c2fo0A_ |
| 1B | BE405749.1 | 4,409,318 | | | | |
| 1B | *TraesAK58CH1B01G008800* | 4,676,665 | | NBS | | disease resistance rpp13-like protein 4, c6j5tC_ |
| 1B | *TraesAK58CH1B01G010100* | 6,010,958 | CC,LRR,NB-ARC | CNL | | disease resistance rpp13-like protein 4, c6j5tC_ |
| 1B | *Ta_Gamma_Sec_19_1RS* | 8,722,719 | | | | |
| 1B | *TraesAK58CH1B01G013200* | 9,385,326 | LRR,STTK,TM | RLK | | lrr receptor-like serine/threonine-protein kinase gso1, c6s6qB |
| 1B | *TraesAK58CH1B01G013600* | 9,484,644 | LRR,STTK,TM | RLK | | lrr receptor-like serine/threonine-protein kinase gso1, c6s6qB |
| 1B | *TraesAK58CH1B01G014500* | 10,083,072 | LRR,STTK,TM | RLK | | lrr receptor-like serine/threonine-protein kinase gso1, c6s6qB |
| 1B | *TraesAK58CH1B01G014720* | 10,187,322 | LRR,STTK,TM | RLK | | lrr receptor-like serine/threonine-protein kinase gso1, c6s6qB |
| 1B | *TraesAK58CH1B01G015820* | 12,022,400 | CC,LRR,NB-ARC | CNL | | disease resistance rpp13-like protein 4, c6j5tC_ |
| 1B | *TraesAK58CH1B01G015800* | 12,221,902 | CC,LRR,NB-ARC | CNL | | tyrosine-protein kinase abl1, c2fo0A_ |
| 1B | *TraesAK58CH1B01G015900* | 12,402,413 | CC,LRR,NB-ARC | CNL | | disease resistance rpp13-like protein 4, c6j5tC_ |
| 1B | BE196644.3 | 15,161,126 | | | | |
| 1B | *TraesAK58CH1B01G021900* | 17,435,279 | LRR,STTK,TM | RLK | | receptor-like protein kinase anxur1, c5y96A_ |
| 1B | *Ta_Omega_Sec_1_1RS* | 18,690,191 | | | | |

**Table S9 RGA gene models analysis on the AK58 genome.**

| Chr | Unique to AK58 | Start | Domain | Class | absent from 1BS CS | domain from Phyre2 analysis, 3D fold for domain |
|---|---|---|---|---|---|---|
| 1B | BE444266.1 | 24,414,912 | | | | |
| 1B | *TraesAK58CH1B01G025900* | 24,625,793 | CC,LRR,NB-ARC,TM | CNL | | disease resistance rpp13-like protein 4, c6j5tC_ |
| 1B | *TraesAK58CH1B01G026600* | 25,432,924 | CC,LRR,NB-ARC | CNL | | disease resistance rpp13-like protein 4, c6j5tC_ |
| 1B | *TraesAK58CH1B01G029200* | 27,920,991 | CC,LRR,NB-ARC | CNL | | disease resistance rpp13-like protein 4, c6j5tC_ |
| 1B | *TraesAK58CH1B01G039510* | 42,207,118 | CC,TM | TM-CC | | nucleoporin gle1, c3peuB_ |
| 1B | *TraesAK58CH1B01G040100* | 42,493,998 | LRR,STTK,TM | RLK | | tyrosine-protein kinase abl1, c2fo0A_ |
| 1B | *TraesAK58CH1B01G081300* | 115,967,305 | LRR,NB-ARC | NL | | disease resistance rpp13-like protein 4, c6j5tC_ |

**green** highlights indicate no cross match with 1BS gene models (based on En

**red** highlights indicate short homologies exists in larger gene families

TraesAK58CH1B01G008800 — disease resistance rpp13-like protein 4, c6j5tC_

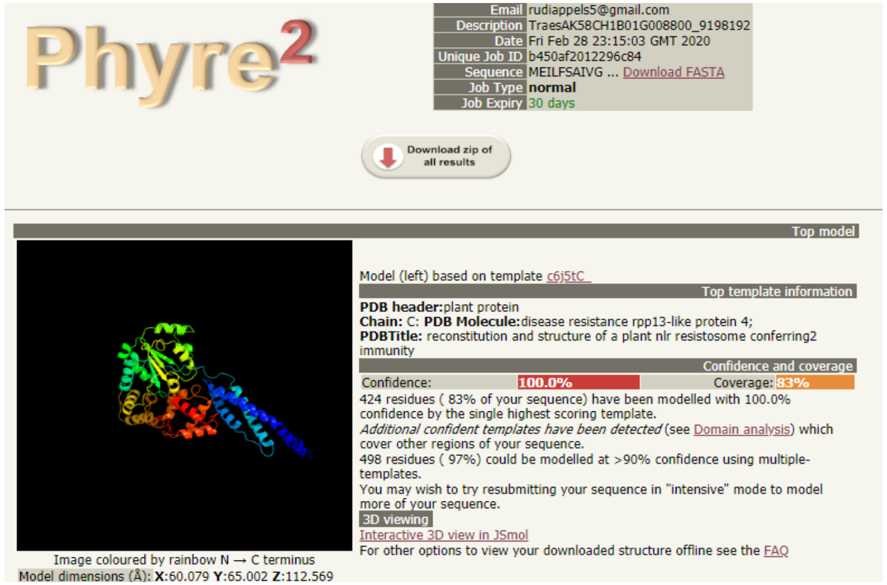

**Table S9 RGA gene models analysis on the AK58 genome.**

| Chr | Unique to AK58 | Start | Domain | Class | absent from 1BS CS | domain from Phyre2 analysis, 3D fold for domain |
|---|---|---|---|---|---|---|
| | TraesAK58CH1B01G010100 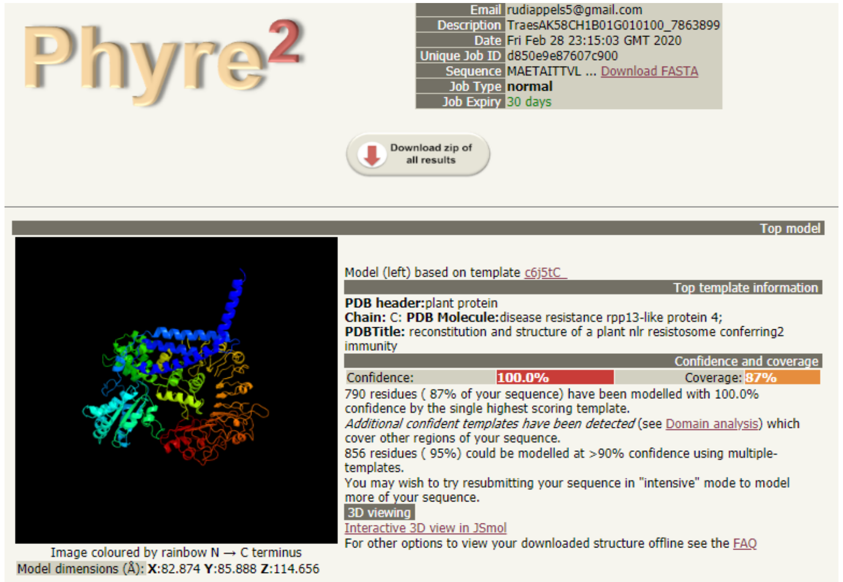 | | | | | disease resistance rpp13-like protein 4, c6j5tC_ |
| | TraesAK58CH1B01G081300 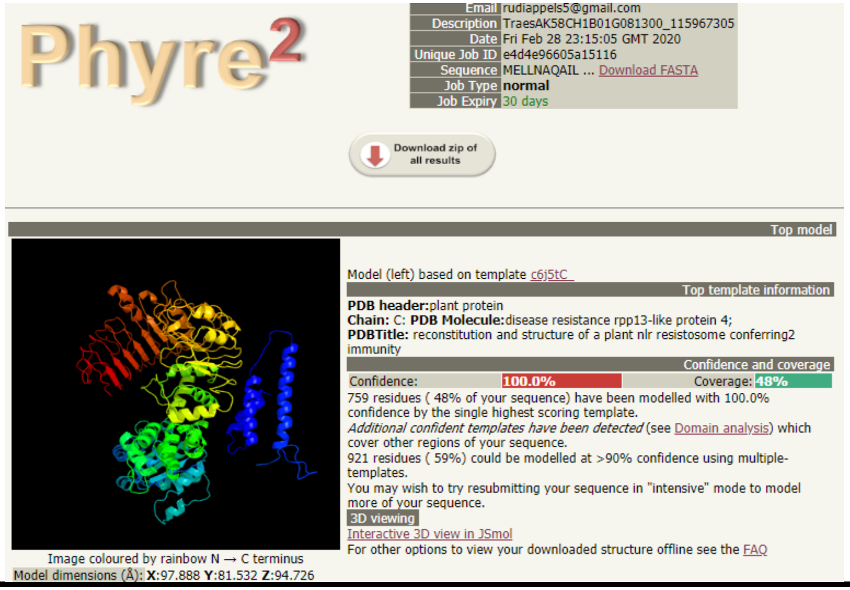 | | | | | disease resistance rpp13-like protein 4, c6j5tC_ |

**Table S9 RGA gene models analysis on the AK58 genome.**

| Chr | Unique to AK58 | Start | Domain | Class | absent from 1BS CS | domain from Phyre2 analysis, 3D fold for domain |
|---|---|---|---|---|---|---|
| | TraesAK58CH1B01G014720 (representative) | | | | | lrr receptor-like serine/threonine-protein kinase gso1; c6s6qB |

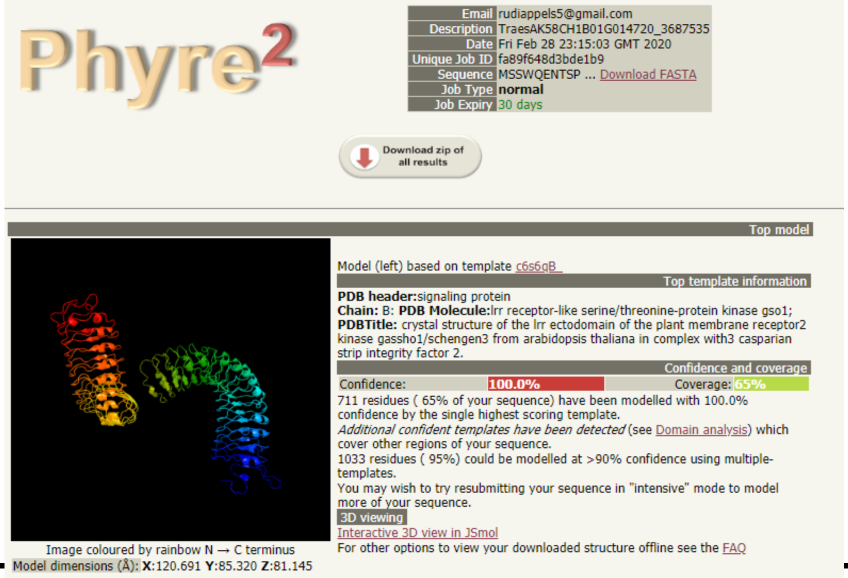
